# Decoding and Targeting Coordinated CDKN1A and CDKN2A Senescence Programs in ECM-Dominant Cardiovascular Pathologies

**DOI:** 10.64898/2026.06.18.732453

**Authors:** Fu Gao, Anthony Fung, Di Zhang, Xing Lou, Jungmin Nam, Mingyu Yang, Xiaolong Tian, Negin Farzad, Dejiang Wang, Gaocai Li, Xiangjun Di, Shujie He, Mei Zhong, Arnar Geirsson, Yang Liu, Rong Fan

**Author notes:** Correspondence should be addressed to (R.F.), (Y.L.), (A.G.).

## Abstract

Cellular senescence is a hallmark of aging and an emerging therapeutic target; however, its role as a context-specific driver of disease remains incompletely defined, and senolytic therapies have shown inconsistent clinical benefit. Here, we identify extracellular matrix (ECM)-dominant pathologies as a major class of senescence-driven disease, characterized by inflammation, matrix degeneration, and progressive tissue dysfunction. Using integrated single-cell transcriptomics, spatial profiling, and multiplex imaging across human specimens and murine models, we demonstrate that senescent fibroblasts, rather than canonical myofibroblasts, constitute the principal disease-driving cell state in myxomatous mitral valve disease (MMVD) and related conditions. These cells exhibit coordinated CDKN1A⁺ inflammatory and CDKN2A⁺ ECM-remodeling programs that form a feed-forward circuit linking immune activation to matrix disorganization and functional decline. Senescence extends beyond fibroblasts to endothelial and immune compartments, establishing a multicellular senescent milieu that reinforces intercellular crosstalk and disease progression. Senolytic treatment (dasatinib plus quercetin or fisetin) restores ECM architecture and improves cardiac function, outperforming pathway-specific anti-inflammatory and antifibrotic approaches. Cross-disease analyses further reveal conservation of this coordinated CDKN1A/CDKN2A senescence programs across multiple ECM-dominant cardiovascular diseases, including aortic aneurysm and calcific valve disease. Notably, in vivo single-cell transcriptomic profiling following multiple senolytic treatments provides whole-transcriptome resolution of context-dependent cellular responses. Collectively, these findings establish context-specific senescence as a central organizing mechanism in ECM-dominant diseases and support a shift from generalized anti-aging strategies toward precision senolytic prevention or therapy. Given that valvular and aortic diseases affect millions and increase markedly with age to a prevalence comparable to major cancers, these results indicate a potential solution to a substantial and underrecognized clinical burden.

## INTRODUCTION

Aging is a major risk factor for cardiovascular disease (CVD) and ECM degeneration, and accumulating evidence supports cellular senescence as a key driver of age-associated tissue dysfunction in these conditions^1–4^. Senescent cells undergo cell-cycle arrest and acquire distinct transcriptional and secretory programs that modulate inflammation, matrix remodeling, and cellular homeostasis^5^. These properties have motivated the development of senolytic therapies, several of which have entered early-phase clinical trials. However, clinical translation has been limited and therapeutic effects modest. For example, in a first-in-human open-label pilot study of dasatinib plus quercetin (D+Q) in idiopathic pulmonary fibrosis (n=14), treatment led to modest improvements in physical measures, including 6-minute walk distance, whereas pulmonary function remained unchanged^6^. In a separate open-label pilot study in diabetic kidney disease (n=9), D+Q reduced senescent cell burden and circulating senescence-associated secretory phenotype (SASP) factors; however, no clear improvement in organ-level efficacy was observed^7^. Clinical data in CVD remain sparse, and ongoing trials have yet to show definitive efficacy. These findings highlight a central mechanistic limitation: current senolytic strategies are largely guided by bulk measurements with limited resolution of the specific cell types and tissue contexts that drive disease. As a result, senescence is often treated as a generalized hallmark of aging rather than a context-dependent, cell-state-specific mechanism. A key unresolved question is whether the senescent phenotype is causative in disease pathogenesis or instead reflects a downstream consequence of disease progression. Relatedly, it remains unclear how senescence-associated programs vary across tissues with distinct structural and functional demands, and whether senolytic therapies primarily suppress downstream disease phenotypes or eliminate upstream senescent cell populations that drive pathology. These gaps underscore the need for single-cell and spatially resolved approached in human tissues to define the precise cellular programs through which senescence contributes to age-associated disease.

Cardiovascular tissues can be conceptualized along a spectrum from cell-dominant to ECM-dominant systems, reflecting whether tissue function is primarily driven by active cellular processes or by the structural and mechanical properties of the ECM^8^. The myocardium represents a prototypical cell-dominant tissue, whereas structures such as cardiac valves and the aorta exemplify ECM-dominant tissues^9,10^. In these tissues, disruption of ECM organization, composition, and biomechanical properties serves as a primary driver of tissue dysfunction, with cellular phenotypes emerging in response to altered microenvironmental cues^10^. ECM-dominant cardiovascular pathologies–including cardiac valve disease and aortic aneurysm–show a marked increase with age^11,12^. Degenerative mitral valve disease affects more than 10% of individuals over 75 years of age^11^, while aortic aneurysms are present in up to 7% of men over 75 years of age^13^, underscoring a substantial and age-dependent clinical burden. In contrast to myocardium, where dysfunction is primarily driven by impaired contractility, pathology in ECM-dominant tissues is fundamentally rooted in matrix disorganization and biomechanical failure^10^. Disease progression in these tissues is characterized by persistent inflammatory signaling, structural matrix remodeling, and impaired resolution of cellular stress responses^14,15^. Although inflammation and fibrosis are well-established features, emerging evidence from multi-omics and single-cell studies suggest that these processes arise from coordinated, cell-state-specific responses to cumulative tissue stress rather than acting as independent drivers. However, the mechanistic relationships linking these processes remain incompletely defined, particularly with respect to cellular senescence and its role in integrating inflammatory and matrix-remodeling programs.

In this context, fibroblast–the principal source of ECM synthesis and key contributors to ECM remodeling–exhibit disease-associated transcriptional programs enriched for inflammatory mediators, (e.g., CXCL and CCL families), ECM components (including collagens, elastin, and glycosaminoglycans), and matrix-modifying enzymes, such as matrix metalloproteinases^16,17^. These coordinated features link immune activation to ECM remodeling and degeneration. Notably, this transcriptional profile is highly consistent with senescence-associated programs; however, senescence is often invoked broadly without precise definition of its tissue- and cell-type-specific roles. p16 (*CDKN2A*) and p21 (*CDKN1A*) are two key cyclin-dependent kinase inhibitors that mediate cell cycle arrest during cellular senescence, with p16 primarily acting through the Rb pathway to enforce stable, irreversible arrest, while p21 functions downstream of p53 to initiate arrest in response to DNA damage. While p16 is a classic marker of irreversible senescence, p21 exhibits a more nuanced, counterintuitive role: single-cell analysis of its early dynamics after non-lethal chemotherapy reveals that senescence-fated cells display lower initial p21 levels than proliferation-fated cells, with modest p21 increases falling into a “Goldilocks zone” that unexpectedly promotes proliferation^18^.

Currently, the prevailing paradigm emphasizes myofibroblast activation as the central driver of fibrotic remodeling^19^. However, this framework does not fully account for the coordinated inflammatory and degenerative ECM changes observed in ECM-dominant pathologies, nor does it explain their strong association with aging in humans. Importantly, senescence is not a uniform state but comprises heterogeneous, context-dependent programs^2,3^, and the specific senescence-associated states that drive disease phenotypes remain incompletely defined. We therefore hypothesized that stable fibroblast states marked by regulators such as *CDKN1A* and *CDKN2A*, consistent with senescence-associated programs, represent functionally distinct cell states that coordinate inflammatory and ECM-remodeling processes. These states may form an integrated pathogenic circuit linking immune activation to maladaptive matrix remodeling, thereby driving disease progression in ECM-dominant conditions.

Here, we investigate ECM-dominant cardiovascular pathologies as a model to define the context-specific role of senescence in disease as well as senotherapeutic responses, using integrated single-cell transcriptomic and spatial analyses across human and murine tissues. We identify senescent fibroblasts, rather than canonical myofibroblasts, as a dominant and previously unrecognized disease-associated cell state in ECM-rich tissues. These cells exhibit distinct but highly coordinated *CDKN1A* inflammatory and *CDKN1A* ECM-remodeling senescence programs that link immune activation to progressive matrix disorganization and tissue dysfunction. In addition, they display immune-evasive and apoptosis-resistant transcriptional features, consistent with persistent pathological activity. Notably, senescence extends beyond fibroblasts to immune compartments, establishing a multicellular senescent milieu that reinforces intercellular crosstalk and disease progression. Functionally, senolytic therapies restore ECM architecture and improve organ-level physiology, as assessed by echocardiography, and outperform pathway-restricted anti-inflammatory and antifibrotic interventions. Importantly, single-cell transcriptomic profiling following senolytic treatment further provides direct in vivo evidence of cell-type-specific responses and mechanistic resolution of therapeutic effects. Moreover, multiplex imaging reveals spatially distinct patterns of immune-associated senescence across anatomical regions within the cardiovascular system.

Together, these findings establish senescence as a context-dependent and mechanistically central driver of ECM-dominant pathologies, rather than a generalized downstream consequence of aging. More broadly, this work provides a framework for the precision deployment of senolytic therapies and offers a mechanistic explanation for the variable outcomes observed in current clinical studies by highlighting the importance of disease-, tissue-, and cell-state-specific senescence programs.

## RESULTS

### Study rationale, design, and workflow

We selected valvular heart disease (VHD)–a prototypical ECM-dominant CVD–as a model system to interrogate the senescence–inflammation–ECM axis. In both aortic and mitral valve disease, pathological ECM remodeling–characterized by fibrosis, collagen disorganization, proteoglycan accumulation, and, in the aortic valve^20^, calcification–leads to leaflet stiffening or degeneration presumably through activation of fibroblast toward myofibroblast- and osteogenic-like states. These changes ultimately result in hemodynamic impairment, ventricular remodeling, and heart failure^21^. Despite a disease burden comparable to that of major cancers, VHD remains relatively understudied. This gap stems in part from a historical focus on end-stage intervention: current treatments are largely surgical or transcatheter replacements applied after irreversible damage has occurred, leaving patients with procedural risks, long-term complications, and compromised quality of life. Consequently, earlier, mechanism-based therapeutic strategies would be ideal but have been limited by an incomplete understanding of VHD pathobiology, particularly in early-stage human disease.

In this work, we initially sought to test a prevailing assumption that myofibroblast activation is the central driver of ECM remodeling and fibrosis in VHD. Unexpectedly, we found that cellular senescence represents a more prominent molecular feature, suggesting a previously unrecognized role in disease pathogenesis. Using a three-stage study design (Figure 1A), we first integrated histological, molecular, and transcriptomic analyses to characterize human MMVD, one of the most common forms of VHD. Rather than observing dominant myofibroblast expansion, we identified a marked enrichment of senescent fibroblasts as a major disease-associated cell population linked to inflammation with ECM disorganization. In addition, senescence-associated immune cell populations contribute to disease progression through coordinated crosstalk with fibroblasts, further supporting a fundamentally distinct pathogenic paradigm. We next performed functional validation by selectively eliminating senescence-associated cells using D+Q or fisetin while in comparison applying anti-inflammatory and antifibrotic interventions to directly compare senescence-targeted and non-senolytic strategies. These treatments were administered in both preventive (prior to disease onset) and interventional (after disease establishment). Finally, we extended our analyses to additional ECM-dominant cardiovascular conditions to evaluate the broader relevance of senescence-driven ECM remodeling as a unifying and therapeutically targetable mechanism across this class of disorders.

**Figure 1.**
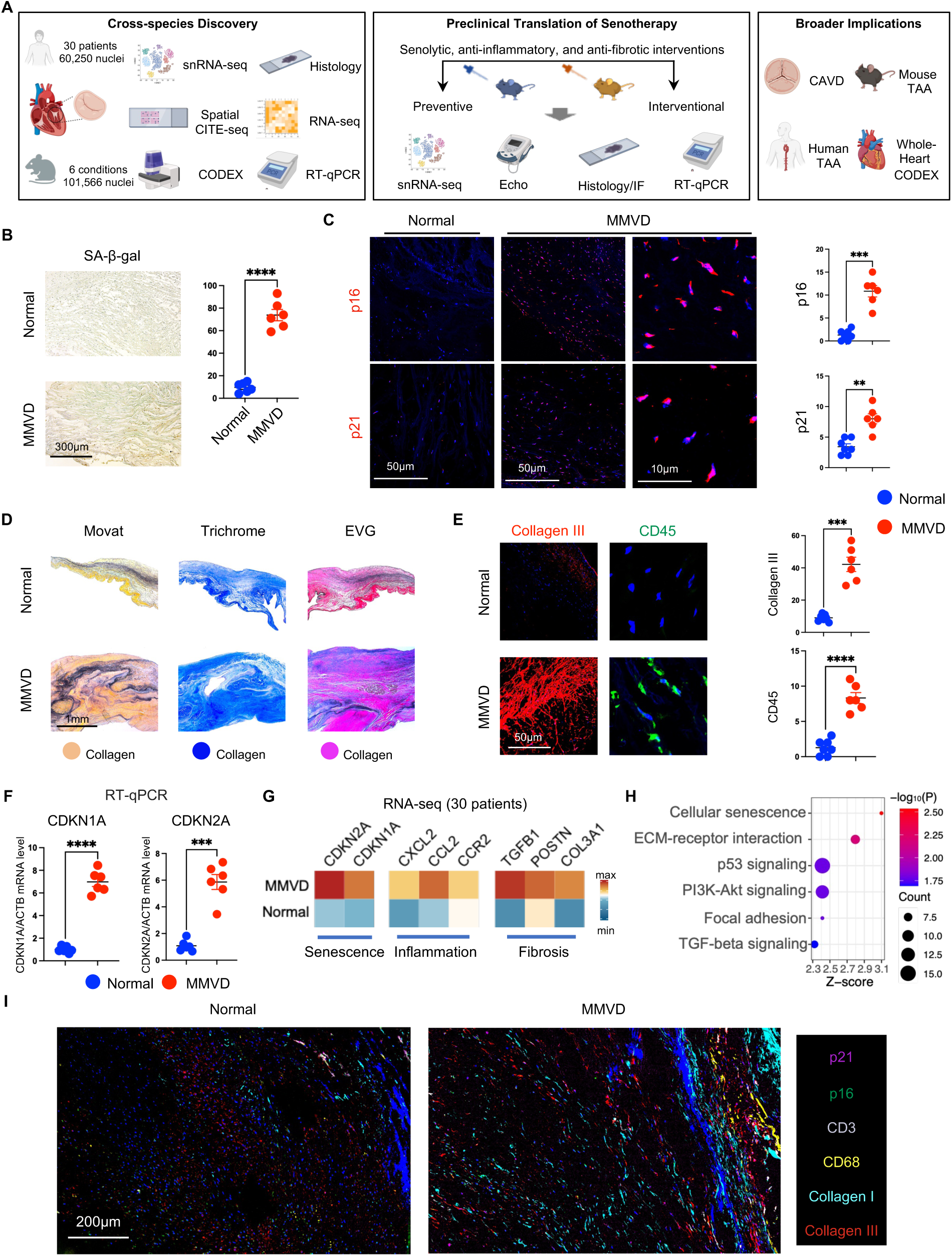
Study design and identification of senescence-associated fibroinflammatory programs in human heart valve disease. **(A)** Overview of the study design. **(B)** Representative images of senescence-associated β-galactosidase (SA-β-gal) staining in normal and myxomatous mitral valve disease (MMVD) tissues. **(C)** Representative immunofluorescence staining of p16 and p21 in mitral valve tissues, with corresponding quantitative analyses. **(D)** Representative histological sections stained with Movat’s pentachrome (Movat), Masson’s trichrome (Trichrome), and elastin van Gieson (EVG), with collagen visualized as yellow in Movat, blue in Trichrome, and red in EVG staining. **(E)** Representative immunofluorescence staining of collagen III and CD45, with corresponding quantitative analyses. **(F)** Scatter plots showing mRNA expression levels of CDKN1A and CDKN2A measured by RT-qPCR. **(G)** Heatmap showing average gene expression levels in normal and MMVD tissues derived from bulk RNA-seq, with values scaled by gene for visualization. **(H)** Bubble plot showing representative enriched KEGG pathways derived from differentially expressed genes (DEGs) between MMVD and normal mitral valve tissues. **(I)** Representative multiplex immunofluorescence (CODEX) images of normal and MMVD mitral valve tissues. Data are presented as mean ± SEM. Statistical significance was determined using unpaired t-tests: ***p<0.001, and ****p<0.0001.

### Senescence-associated fibroinflammatory remodeling in patients

Given the strong age dependence of VHD, we first assessed senescence burden in human mitral valve tissues. Senescence-associated β-galactosidase (SA-β-gal) activity was increased in MMVD valves compared with normal controls (Figure 1B). This increase was corroborated by immunofluorescence staining, which revealed significantly elevated expression of canonical senescence markers, including p21 and p16, indisease tissues (Figure 1C), consistent with accumulation of senescence-associated cells in diseased valves. Consistent with these findings, histopathological analyses using Movat pentachrome, Masson’s trichrome, and elastin van Gieson (EVG) staining demonstrated pronounced ECM disorganization and fibrotic remodeling in MMVD valves (Figure 1D). Collagen content was significantly increased, which was further validated by collagen III staining (Figure 1E). Notably, regions of excessive collagen deposition were accompanied by prominent immune cell infiltration, consistent with a fibrotically remodeled and inflamed valvular microenvironment^14,22,23^ (Figure 1E).

To define the molecular programs underlying these pathological changes, we profiled transcriptional alterations at both targeted and global levels. Reverse transcription quantitative polymerase chain reaction (RT-qPCR) confirmed significant upregulation of genes associated with senescence (*CDKN2A* and *CDKN1A*), inflammation (*CCL2* and *IL6*), and fibrosis (*TGFB1, COL1A1, COL3A1*) in MMVD tissues relative to controls (Figure 1F and S1A). Extending these findings, bulk RNA sequencing of human mitral valves from control donors (n = 10) and MMVD patients (n = 20) (Table S1) revealed broad activation of senescent, inflammatory, and fibrotic gene expression programs in diseased valves (Figure 1G). Pathway enrichment analyses further supported robust activation of senescence-, inflammation-, and ECM-related signaling pathways in MMVD (Figure 1H). Finally, high-dimensional multiplexed immunofluorescence imaging (CODEX) enabled spatial validation of these signatures at single-cell resolution^24^. This analysis demonstrated coordinated spatial accumulation of senescence-associated cells, immune infiltrates, and profibrotic ECM remodeling within MMVD valves (Figure 1I and S2A), supporting the presence of a structured senescent and fibroinflammatory microenvironment in diseased tissue.

Together, these complementary analyses establish that human heart mitral valve degeneration is likely associated with a coordinated senescence-associated fibroinflammatory microenvironment tightly coupled to ECM disorganization and fibrosis. Beyond defining a key feature of valvular pathology, these findings suggest that cellular senescence may represent a fundamental and previously underappreciated driver of ECM remodeling in human disease.

### Distinct CDKN1A-high and CDKN2A-high fibroblast states cooperatively drive pathological ECM remodeling

To dissect the cellular mechanisms underlying the senescence-immune-ECM axis and to define the cell types contributing to senescence-associated programs, we performed snRNA-seq of human mitral valve tissues from control donors (n = 10) and MMVD patients (n = 12) (Table S2). Nuclei isolation from mitral valves is technically challenging due to dense ECM deposition and low intrinsic cellularity. After stringent quality control, we obtained 60,250 high-quality nuclei from 12 patients and 10 controls. Unsupervised clustering identified five major cell populations (Figure 2A and S3A), which were annotated using established lineage-specific marker genes^25,26^ (Figure 2B). The disease valves exhibited a relative expansion of macrophages compared with control valves (Figure S3B), consistent with the inflammatory microenvironment observed histologically. In parallel, spatial-CITE-seq revealed enrichment of senescence-, immune-, and fibrosis-associated RNA and protein expression in MMVD tissues, indicating coordinated activation of these pathological processes in situ^27,28^ (Figure 2C and Table S3).

**Figure 2.**
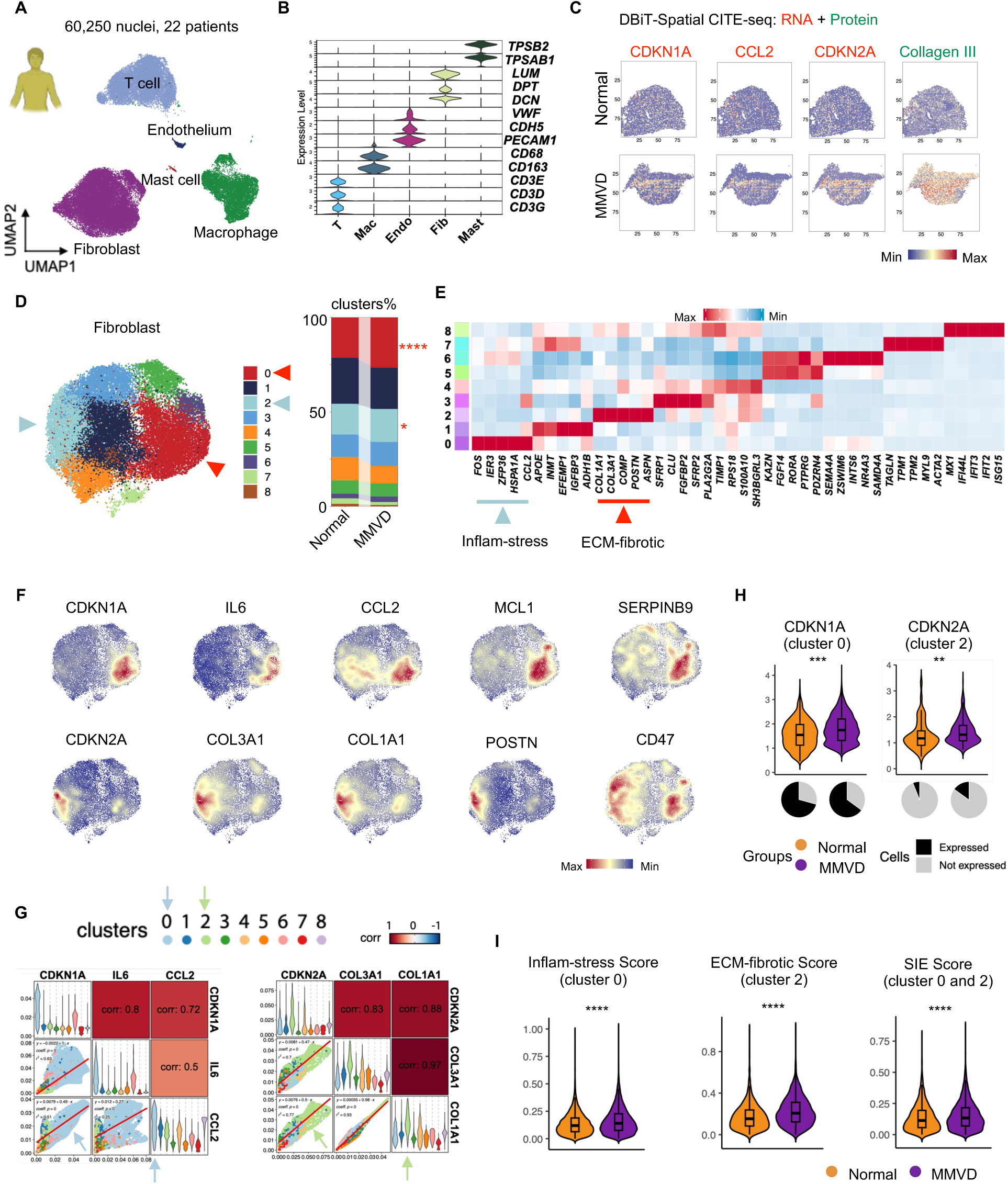
Distinct functional states of senescence-associated fibroblasts in myxomatous mitral valve disease. **(A)** UMAP showing the major cellular components of normal and myxomatous mitral valve tissues. **(B)** Violin plots showing expression of canonical marker genes across major cell types. Endothelial cells (Endo), fibroblasts (Fib), macrophages (Mac), T cells (T). **(C)** Spatial-CITE-seq maps showing the spatial distribution of representative senescence and inflammatory markers (CDKN1A, CDKN2A, CCL2; RNA) and the fibrotic marker collagen III (protein) in normal and myxomatous mitral valves. **(D)** UMAP of unsupervised fibroblast clustering, with arrows indicating fibroblast clusters significantly expanded in MMVD and enriched for CDKN1A or CDKN2A; stacked bar plots show the proportional distribution of fibroblast clusters in normal and MMVD samples. **(E)** Heatmap showing the top five marker genes for each fibroblast cluster, with arrows highlighting clusters enriched for CDKN2A and CDKN1A, corresponding to ECM-fibrotic and inflammatory-stress (Inflam-stress) programs, respectively. **(F)** Expression of senescence (CDKN1A, CDKN2A), inflammatory (IL6, CCL2), immune-evasion/survival (MCL1, SERPINB9, CD47), and fibrotic (COL3A1, COL1A1, POSTN) genes in fibroblasts. **(G)** Correlation analysis showing strong associations between CDKN1A, IL6, and CCL2 predominantly in cluster 0 (left), and between CDKN2A, COL3A1, and COL1A1 predominantly in cluster 2 (right) across fibroblast clusters. **(H)** Violin plots showing CDKN1A expression in cluster 0 and CDKN2A expression in cluster 2 in normal and MMVD samples, with pie charts below depicting the proportions of cells expressing or not expressing each gene in the corresponding groups. **(I)** Violin plots comparing inflammatory-stress, ECM-fibrotic, and Survival and Immune-Evasion (SIE) scores between normal and MMVD fibroblast clusters. Data are presented as mean ± SEM. Statistical significance was determined using Wilcoxon tests: **P < 0.01, ***P < 0.001, and ****P < 0.0001.

We next focused on fibroblasts, the predominant cell population responsible for maintaining ECM-dominant tissue structure, mechanical integrity, and matrix homeostasis. Unsupervised clustering identified substantial fibroblast heterogeneity^29–31^ (Figure 2D). Among nine fibroblast clusters, clusters 0 and 2 were selectively expanded in disease relative to controls. Furthermore, unsupervised cluster-enriched top marker genes revealed distinct fibroblast transcriptional states. Specifically, cluster 0 was characterized by a stress-activated inflammatory transcriptional program, whereas cluster 2 exhibited an ECM-remodeling, pro-fibrotic program (Figure 2E).

We were curious about the involvement of senescence and related programs in in these fibroblast populations and thus examined representative senescence, inflammatory, and ECM/fibrotic markers. Feature plots demonstrated that cluster 0 exhibited elevated expression of *CDKN1A,* a gene encoding p21, a crucial cyclin-dependent kinase inhibitor that drives cell cycle arrest in senescence, together with inflammatory and stress-responsive genes, including *IL6* and *CCL2*. In contrast, cluster 2 showed preferential enrichment of *CDKN2A*, which encodes the protein p16INK4a, a master regulator of cellular senescence, and triggers irreversible cell cycle arrest in response to stress or aging, along with fibrotic ECM genes such as *COL1A1, COL3A1*, and *POSTN* (Figure 2F). Correlation analyses revealed strong positive associations between *CDKN1A* and inflammatory gene expression within cluster 0, whereas *CDKN2A* expression in cluster 2 was tightly correlated with ECM-remodeling and profibrotic programs; these associations were weaker in other fibroblast clusters (Figure 2G). Notably, canonical myofibroblast markers (*ACTA2* and *MYH11*) were enriched in a distinct fibroblast cluster and showed minimal overlap with the senescent-like populations (Figure S3C). Quantitative analyses further showed that *CDKN1A* expression in cluster 0 and *CDKN2A* expression in cluster 2 were significantly increased in diseased valves compared with controls, accompanied by a higher proportion of cells expressing these two senescence signature genes within each respective cluster (Figure 2H). Consistently, an inflammation-stress score was significantly elevated in cluster 0, whereas an ECM-fibrotic remodeling score was significantly increased in cluster 2 in MMVD tissues (Figure 2I and Tables S4 and S5).

Under physiological conditions, senescent or damaged cells are subject to immune surveillance and clearance^5,32,33^; however, this process appears to be compromised in diseased valves. Consistent with this notion, the SIE (Survival and Immune-Evasion) score was significantly increased in diseased compared with control valves in both clusters 0 and 2, supporting enhanced senescent cell persistence and immune evasion^5,33–36^ (Figure 2I and Table S6). Notably, these two fibroblast clusters exhibited distinct immune-protective programs. In cluster 0, *CDKN1A* enrichment was associated with increased expression of *MCL1*, an anti-apoptotic *BCL2* family member that supports survival under stress conditions, together with *SERPINB9*, a granzyme B inhibitor conferring resistance to cytotoxic T cell-mediated killing^37^. In contrast, the *CDKN2A*-enriched cluster 2 showed an association with *CD47*, a canonical “don’t-eat-me” signal that suppresses macrophage-mediated phagocytosis (Figure 2F and S3D)^38–40^. These findings indicate that *CDKN1A*-high and *CDKN2A*-high fibroblast states deploy complementary immune-evasion strategies that may promote persistence of senescence-associated cells in disease.

Together, these data demonstrate that among multiple fibroblast subpopulations, *CDKN1A*-high inflammatory and *CDKN2A*-high profibrotic fibroblast states are selectively expanded in MMVD and are associated with pathological ECM remodeling and senescence-linked cell persistence.

### Senescence-associated transcriptional features of immune cell populations

We next examined the immune compartment to determine whether senescence-associated transcriptional programs extend beyond fibroblasts and shape disease-associated immune states. In T cells, unsupervised clustering identified eight distinct clusters (Figure 3A and Figure S4A). Among these, clusters 0 and 3 were significantly expanded in the diseased samples, whereas cluster 2 was significantly reduced compared with controls (Figure 3A), indicating disease-associated remodeling of the T cell landscape^41–43^.

**Figure 3.**
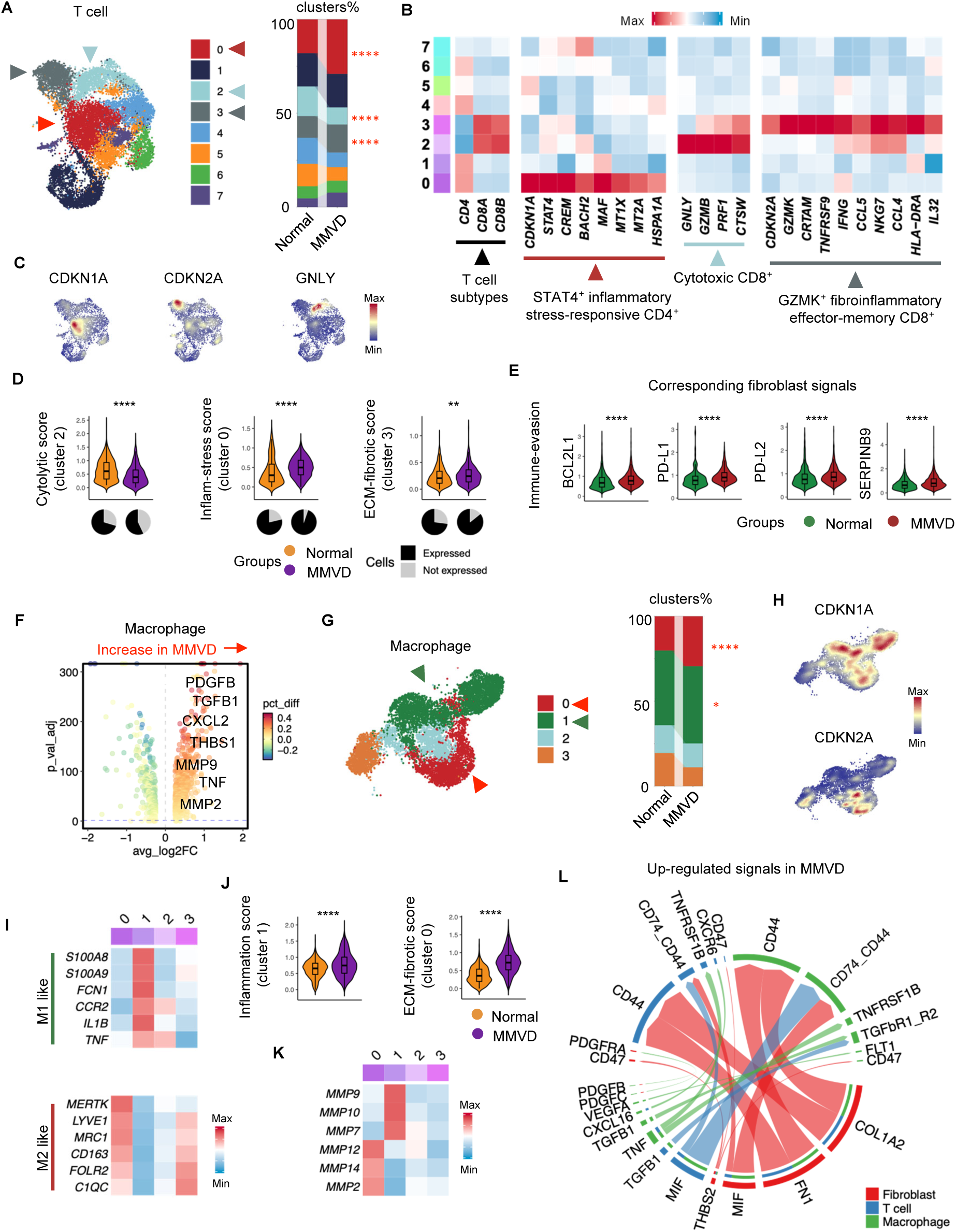
Senescence-associated immune cell states in human mitral valve disease. **(A)** UMAP of unsupervised T cell clustering, with arrows indicating T cell clusters whose proportions are significantly altered between normal and MMVD samples and are enriched for CDKN2A, CDKN1A, and GNLY expression; stacked bar plots show the proportional distribution of T cell clusters in normal and MMVD samples. **(B)** Heatmap showing representative marker genes used for annotation of T cell clusters, with arrows indicating major T cell subtypes (CD4⁺ and CD8⁺) and distinct functional states, including cluster 0 (CDKN1A⁺) identified as STAT4⁺ inflammatory stress-responsive CD4⁺ T cells, cluster 3 (CDKN2A⁺) identified as GZMK⁺ fibroinflammatory effector-memory CD8⁺ T cells, and cluster 2 (GNLY⁺) identified as cytotoxic CD8⁺ T cells. **(C)** Expression of CDKN1A, CDKN2A, and GNLY across T cells. **(D)** Violin plots comparing cytolytic, inflammatory-stress, and ECM-fibrotic scores across T cell clusters in normal and MMVD samples, with pie charts indicating the proportions of cells expressing or not expressing the corresponding gene sets in each group. **(E)** Violin plots showing expression of immune-evasion-associated genes in fibroblasts from normal and MMVD samples. **(F)** DEGs between macrophages from normal and MMVD samples, with representative genes upregulated in MMVD highlighted. **(G)** UMAP of unsupervised macrophage clustering, with arrows indicating macrophage clusters significantly expanded in MMVD and enriched for CDKN2A and CDKN1A expression; stacked bar plots show the proportional distribution of macrophage clusters in normal and MMVD samples. **(H)** Expression of CDKN1A and CDKN2A in macrophages. **(I)** Heatmap showing representative marker genes defining M1- and M2-like macrophage subtypes. **(J)** Violin plots comparing inflammatory-stress and ECM-fibrotic scores across macrophage clusters in normal and MMVD samples. **(K)** Heatmap showing expression of representative matrix metalloproteinase genes across macrophage clusters. **(L)** Representative ligand-receptor interactions upregulated in MMVD compared with normal mitral valves, with edge thickness reflecting inferred interaction strength. Data are presented as mean ± SEM. Statistical significance was determined using Wilcoxon tests: *P < 0.05, **P < 0.01, and ****P < 0.0001.

Cluster-enriched marker analysis revealed that cluster 0 corresponded to a *STAT4*⁺ inflammatory, stress-responsive CD4⁺ T cell program, characterized by elevated expression of cytokine- associated transcriptional regulators (*STAT4, CREM, MAF, BACH2*) together with oxidative and cellular stress-response genes (*MT1X, MT2A*)^44,45^. Cluster 2 was defined by a cytotoxic CD8⁺ T cell program, marked by high expression of canonical cytolytic molecules (*GNLY, GZMB, PRF1, CTSW*)^46–50^. In contrast, cluster 3 exhibited a *GZMK*⁺ fibroinflammatory effector-memory CD8⁺ T cell program, characterized by expression of inflammatory effector genes (*GZMK, IFNG, IL32*), chemokines (*CCL4, CCL5*), and sustained activation receptors (*TNFRSF9, CRTAM*), together with activation-associated markers (*HLA-DRA, NKG7*). This transcriptional profile is consistent with a fibroinflammatory immune state and suggest a potential role for these T cells in modulating fibroblast activation and ECM remodeling through chronic immune-stromal crosstalk^51–56^ (Figure 3B). Consistent with these annotations, feature plots demonstrated preferential enrichment of *CDKN1A* in the *STAT4*⁺ cluster and *CDKN2A* in the *GZMK*⁺ cluster, whereas expression of cytotoxic effector genes such as *GNLY* was largely restricted to the cytotoxic CD8⁺ cluster (Figure 3C).

Furthermore, relative to normal samples, *STAT4*⁺ cluster 0 T cells in MMVD exhibited increased expression of cytokine- and stress-response genes, including *IFNGR1, CREM,* and *STAT4*, whereas *GZMK*⁺ cluster 3 T cells showed upregulation of genes linked to chronic inflammatory activation and fibroinflammatory signaling potential, including *TGFB1, MIF, TNFRSF1B*, and *CXCR6*. In contrast, cytotoxic CD8⁺ cluster 2 T cells displayed reduced expression of canonical cytotoxic effector genes, including *GNLY, GZMB, and CTSW* in MMVD compared with controls (Figure S4B). To quantitatively assess functional differences among these T-cell states, we applied gene set-based scoring to evaluate cytolytic activity, inflammatory stress responses, and profibrotic potential. Consistent with reduced abundance of cytotoxic CD8 T cells, cytolytic scores in cluster 2 were significantly decreased in disease compared with control^57^ (Figure 3D and Table S7). In contrast, inflammation-stress scores were significantly elevated in cluster 0 (Figure 3D and Table S8), whereas profibrotic scores were significantly increased in cluster 3 in disease (Figure 3D and Table S9). Together, these results support a shift from cytotoxic immune surveillance toward coordinated inflammatory and chronically profibrotic T cell programs in disease. Given the reduction in cytotoxic T cell activity, we next examined whether fibroblasts exhibit altered susceptibility to immune-mediated clearance by assessing genes associated with resistance to T cell cytotoxicity. Compared with controls, fibroblasts from MMVD samples showed significantly increased expression of the anti-apoptotic gene *BCL2L1*, immune checkpoint ligands *PD-L1* (*CD274*) and *PD-L2* (*PDCD1LG2*), and *SERPINB9*, an inhibitor of granzyme B-mediated cytotoxicity (Figure 3E). These findings indicate that diseased fibroblasts acquire an immune-evasive and apoptosis-resistant transcriptional program consistent with reduced susceptibility to cytotoxic T cell-mediated killing^58,59^.

We next investigated the effect in macrophages. Differential expression analysis revealed that macrophages from diseased samples upregulated inflammatory and profibrotic genes, including *PDGFB, TGFB1, THBS1, MMP2, CXCL2,* and *TNF*, consistent with an activated macrophage state with the potential to promote fibroblast activation (Figure 3F). Unsupervised clustering identified four macrophage subpopulations (Figure 3G and Figure S4C). Among these, cluster 0 showed preferential enrichment of *CDKN2A*, whereas cluster 1 was enriched for *CDKN1A* (Figure 3H). Cluster-enriched marker gene analysis indicated that the *CDKN2A*-enriched macrophage cluster exhibited features consistent with a resident, M2-like macrophage program associated with tissue remodeling and fibrosis, whereas the *CDKN1A*-enriched cluster corresponded to a *CCR2*⁺, M1-like monocyte-derived inflammatory macrophage state (Figure 3I), again, indicating distinct yet coordinated *CDKN1A/CDKN2A* senescent cell states. To quantify functional differences, we applied gene set-based scoring for inflammatory and profibrotic activity. Consistent with its inflammatory identity, the *CCR2*⁺ *CDKN1A*-enriched macrophage cluster displayed significantly elevated inflammation scores in MMVD compared with controls^14^ (Figure 3J and Table S4). In contrast, the *CDKN2A*-enriched resident macrophage cluster exhibited significantly higher profibrotic scores in MMVD^60–62^ (Figure 3J and Table S5).

Analysis of ECM-remodeling genes revealed that both macrophage subsets expressed matrix metalloproteinases (MMPs), but with distinct patterns (Figure 3K). The *CCR2*⁺ *CDKN1A*-enriched macrophage cluster preferentially expressed MMPs commonly associated with inflammatory matrix turnover, including *MMP7*, *MMP9*, and *MMP10*^63–65^, whereas the resident M2-like macrophage cluster enriched for *CDKN1A* exhibited higher expression of *MMP2, MMP12*, and *MMP14*, which are more frequently linked to chronic ECM remodeling and fibrotic tissue restructuring^66–68^.

Finally, to delineate intercellular communication networks underlying senescence-associated remodeling, we performed ligand-receptor interaction analysis^69^ (Figure 3L). MMVD samples exhibited enhanced signaling among fibroblasts, T cells, and macrophages, with fibroblasts emerging as a central communication hub. Fibroblast-derived ligands involved in ECM remodeling and immune modulation, including *FN1, THBS2,* and *MIF*, showed increased interactions with immune receptors such as *CD44, CD47,* and *CD74*. Conversely, immune-derived ligands including *TGFB1, PDGFB, TNF,* and *CXCL16* were prominently enriched, consistent with activation of pathways governing inflammation, cell survival, and matrix remodeling.

Together, these findings again indicate the role of distinct and coordinated *CDKN1A* and *CDKN2A* senescence-associated transcriptional programs beyond fibroblasts to establish a multicellular senescent milieu that amplifies fibroinflammatory crosstalk and defines distinct immune states in ECM-dominant disease while fibroblast senescence acts as the central hub, supporting a senescence-associated immune-ECM remodeling axis underlying progressive ECM degeneration.

### The fibrillin-1-deficient mgR mouse model recapitulates senescence-associated features

Fibrillin-1 deficiency leads to a wide range of ECM-associated pathologies in mice including severe myxomatous degeneration of the mitral valve, characterized by ECM disruption and fibrotic remodeling. To determine whether senescence-associated programs observed in human MMVD are recapitulated in this context, we examined the *Fbn1^mgR/mgR^*(mgR) mouse model^70^. We analyzed 12-week-old homozygous mgR mice, with *Fbn1^+/+^* (WT) littermates as controls. Gross examination revealed markedly enlarged mitral valve leaflets in mgR mice (Figure 4A), and histology confirmed pronounced structural alterations, including increased leaflet thickness and architectural disorganization (Figure 4A).

**Figure 4.**
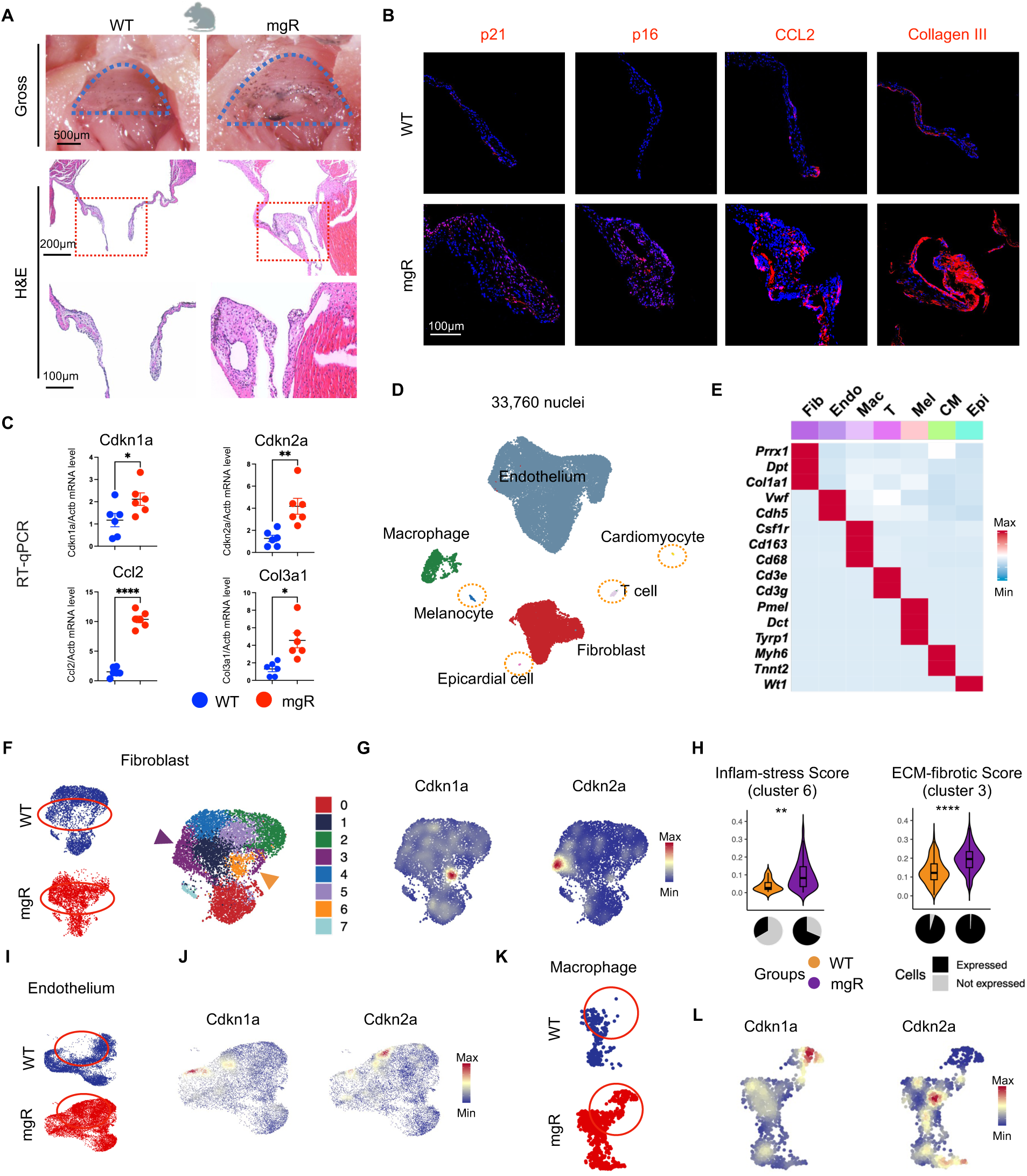
Cellular senescence in the mgR mouse model of mitral valve disease. **(A)** Gross morphology (top) and histological features (bottom) of the mitral valve in wild-type (WT) and mgR mice. **(B)** Representative immunofluorescence staining of p21, p16, CCL2, and collagen III in mitral valve sections. **(C)** Scatter plots showing mRNA expression levels of Cdkn1a, Cdkn2a, Ccl2, and Col3a1 measured by RT-qPCR. **(D)** UMAP showing the cellular composition of WT and mgR mitral valves. **(E)** Heatmap showing the expression of canonical marker genes across major cell types. Cardiomyocyte (CM), epicardial cell (Epi), melanocyte (Mel). **(F)** UMAP of WT and mgR fibroblasts (left) and unsupervised clustering of fibroblasts (right). Arrows indicate fibroblast clusters enriched for Cdkn1a and Cdkn2a expression. **(G)** Expression of Cdkn1a and Cdkn2a in fibroblasts. **(H)** Violin plots comparing transcriptional program scores between WT and mgR fibroblast clusters. **(I)** UMAP of WT and mgR endothelial cells. **(J)** Expression of Cdkn1a and Cdkn2a across endothelial cell clusters. **(K)** UMAP of WT and mgR macrophages. **(L)** Expression of Cdkn1a and Cdkn2a in macrophages. Data are presented as mean ± SEM. Statistical significance was determined using Wilcoxon tests: *P < 0.05, **P < 0.01, and ****P < 0.0001.

Consistent with the human MMVD phenotype, immunofluorescence staining demonstrated increased abundance of p21- and p16-positive cells in mgR mitral valves, accompanied by elevated expression of the inflammatory chemokine CCL2 and excessive collagen III deposition^14,15^ (Figure 4B). RT-qPCR further confirmed increased expression of *Cdkn1a, Cdkn2a, Ccl2*, and *Col3a1* in mgR valves relative to WT controls (Figure 4C). We next performed snRNA-seq on mitral valves from mgR mice and WT littermates. Despite the minute tissue size of murine mitral valves, we captured 33,760 high-quality nuclei across both groups. Unsupervised clustering identified seven major cell populations, including fibroblasts, endothelial cells, macrophages, T cells, cardiomyocytes, epicardial cells, and a small melanocyte population unique to murine valves, based on established lineage marker genes^25,26^ (Figure 4D, 4E and Figure S5A).

We first examined fibroblasts, the predominant stromal population in the mitral valve. Unsupervised clustering identified eight transcriptionally distinct fibroblast subpopulations (Figure 4F and Figure S5B). Two fibroblast clusters (clusters 3 and 6) were selectively expanded in mgR valves (Figure 4F). Feature plots demonstrated enrichment of *Cdkn1a* in cluster 6 and *Cdkn2a* in cluster 3 (Figure 4G). Cluster 6 exhibited increased expression of inflammatory and stress-responsive genes, whereas cluster 3 was enriched for ECM-associated and profibrotic genes (Figure S5C). Consistent with these programs, inflammatory-stress scores were significantly elevated in cluster 6 and ECM-fibrotic scores were increased in cluster 3 in mgR valves relative to WT controls (Figure 4H and Tables S4 and S5).

Given their anatomical position at the blood-tissue interface and their role as a barrier between stromal cells and circulating immune cells, we next examined endothelial cells. Unsupervised clustering identified multiple transcriptionally distinct endothelial subpopulations (Figure 4I and Figure S5D). Two endothelial subpopulations (clusters 3 and 7) were markedly expanded in mgR valves, whereas only a small fraction of WT endothelial cells occupied these states (Figure 4I). Feature plots showed preferential enrichment of *Cdkn1a* in cluster 7 and *Cdkn2a* in cluster 3 (Figure 4J). Clusters 3 and 7 preferentially expressed fibroinflammatory genes, consistent with an activated endothelial program associated with enhanced immune cell adhesion and recruitment, increased endothelial-ECM interactions, and paracrine profibrotic signaling (Figure S5E)^71–77^. Together, these data support senescence-associated endothelial dysfunction and reprogramming in the mgR model. Unsupervised clustering of macrophages identified three subpopulations (Figure 4K and Figure S5F). The macrophage population expanded in mgR valves showed enrichment of both *Cdkn1a* and *Cdkn2a* (Figure 4L). Differential expression analysis identified upregulation of inflammatory and fibrotic mediators, including *Il1b, Fn1, Vcan, Ccr2,* and *Mmp12* in mgR mice (Figure S5G), consistent with a macrophage state integrating inflammatory activation with active ECM remodeling. Thus, the mgR murine model also demonstrated the implication of distinct *Cdkn1a* and *Cdkn2a* senescence programs in ECM-dominant disease phenotype.

Finally, ligand-receptor interaction analysis revealed enhanced intercellular communication among fibroblasts, endothelial cells, and macrophages in mgR valves (Figure S5H). Upregulated interactions were enriched for integrin- and growth factor-mediated pathways, linking fibroblast-derived ECM ligands to receptors on endothelial cells and macrophages. Conversely, inflammatory ligands from immune and endothelial cells engaged fibroblast receptors involved in cell survival, activation, and matrix remodeling. Together, these interactions delineate a senescence-associated communication network consistent with coordinated inflammatory and fibrotic remodeling across cell types in mgR mice.

### Senotherapies prevent and ameliorate valvular heart disease and restore cardiac function in the mouse model

To assess the potential therapeutic targeting of senescence-associated programs and compare the effect against anti-inflammation and anti-fibrosis treatment, we treated mgR mice for 8 weeks (from 4 to 12 weeks of age) with senolytics (D+Q or fisetin), anti-inflammatory agents (RS504393 (CCR2 inhibition; CCR2i) or anti-IL-1β)^15,78^; or antifibrotic agent pirfenidone (PFD)^79^. Gross inspection and histological analyses showed that D+Q, fisetin, CCR2i, and pirfenidone attenuated mitral valve degeneration, whereas anti-IL-1β did not confer appreciable structural improvement (Figure 5A). Quantitative morphometric analyses revealed significant reductions in mitral valve leaflet area and thickness following D+Q, fisetin, CCR2i, or pirfenidone compared with untreated mgR controls (Figure 5B). Among these interventions, senolytic treatments were associated with the most pronounced structural improvement, with valve morphology partially approaching that of WT controls, whereas CCR2 inhibition and pirfenidone showed intermediate effects. Movat pentachrome staining further demonstrated improved ECM organization and decreased collagen accumulation in senolytic-treated mice (Figure 5B). Functional assessment by echocardiography showed that D+Q, fisetin, and CCR2i reduced mitral regurgitation (Figure 5C). By comparison, pirfenidone conferred a weaker functional benefit, consistent with the more limited structural rescue observed histologically. At the cellular level, immunofluorescence analysis showed that senolytic treatments reduced senescence marker-positive cells in the mitral valve, as evidenced by decreased numbers of both p16^+^ and p21^+^ cells (Figure 5D). In addition, D+Q and fisetin reduced the abundance of CCL2⁺ cells. Fibrotic remodeling was further assessed by collagen III staining, which demonstrated reduced pathological ECM deposition following senolytic treatment (Figure 5D).

**Figure 5.**
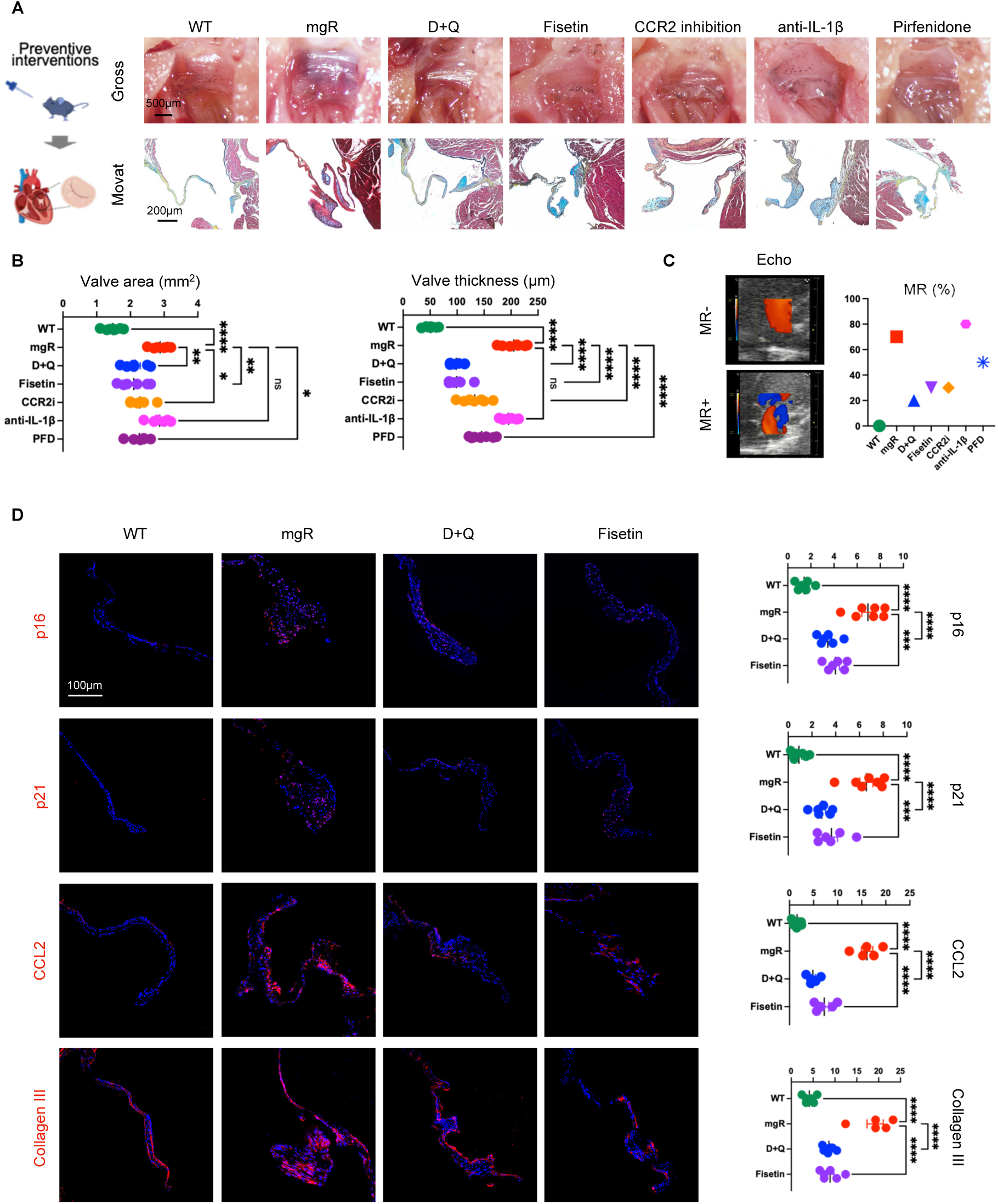
Senolytic therapies ameliorate mitral valve disease and restore cardiac function in mice. **(A)** Gross morphology (top) and histological features (bottom) of the mitral valve across seven experimental groups. **(B)** Quantification of mitral valve area and leaflet thickness across seven experimental groups. CCR2 inhibition (CCR2i); pirfenidone (PFD). **(C)** Representative echocardiographic (Echo) images illustrating normal valve function and mitral regurgitation (MR) (left), and the incidence of MR across the indicated experimental groups (right). **(D)** Representative immunofluorescence staining of p16, p21, CCL2, and collagen III, with corresponding quantitative analyses. Data are shown as individual values with mean ± SEM; ns, not significant; *P < 0.05, **P < 0.01, ***P < 0.001, and ****P < 0.0001 by one-way ANOVA.

In the mgR mouse model, ECM disruption and fibrotic remodeling emerge by one month of age and progressively worsen, leading to multi-organ fibrotic pathologies including mitral valve dysfunction. To assess whether senolytic therapy can ameliorate established disease, mgR mice were treated from 8 to 20 weeks of age with senolytics (D+Q or fisetin), CCR2 inhibition, or pirfenidone. While CCR2 inhibition and pirfenidone produced modest effects, only senolytic treatment significantly reduced mitral valve leaflet area and thickness compared with untreated mgR mice (Figure S6A and S6B). Histological analysis demonstrated restoration of ECM organization with reduced ECM degeneration and fibrotic accumulation, and echocardiography confirmed decreased mitral regurgitation severity following senolytic therapy, with non-senolytic interventions showing limited functional improvement (Figure S6C). Collectively, these findings indicate that senolytic therapy attenuates established leaflet remodeling and improves mitral valve function in MMVD.

### snRNA-seq reveals the necessity to suppress both *Cdkn1a+* and *Cdkn2a+* senescent populations with senolytics to achieve therapeutic efficacy

To determine how distinct therapeutic interventions modulate senescence-associated cellular programs at single-cell resolution, we performed snRNA-seq on mitral valves from six experimental groups: WT, mgR, D+Q, fisetin, CCR2i, and pirfenidone treated mice. After stringent quality control, 101,566 high-quality nuclei were retained for downstream analyses (Figure 6A). Unsupervised clustering identified seven major cell types, including endothelial cells, fibroblasts, macrophages, T cells, cardiomyocytes, melanocytes, and smooth muscle cells (SMC), annotated using established lineage markers (Figure 6A and 6B and Figure S7A).

**Figure 6.**
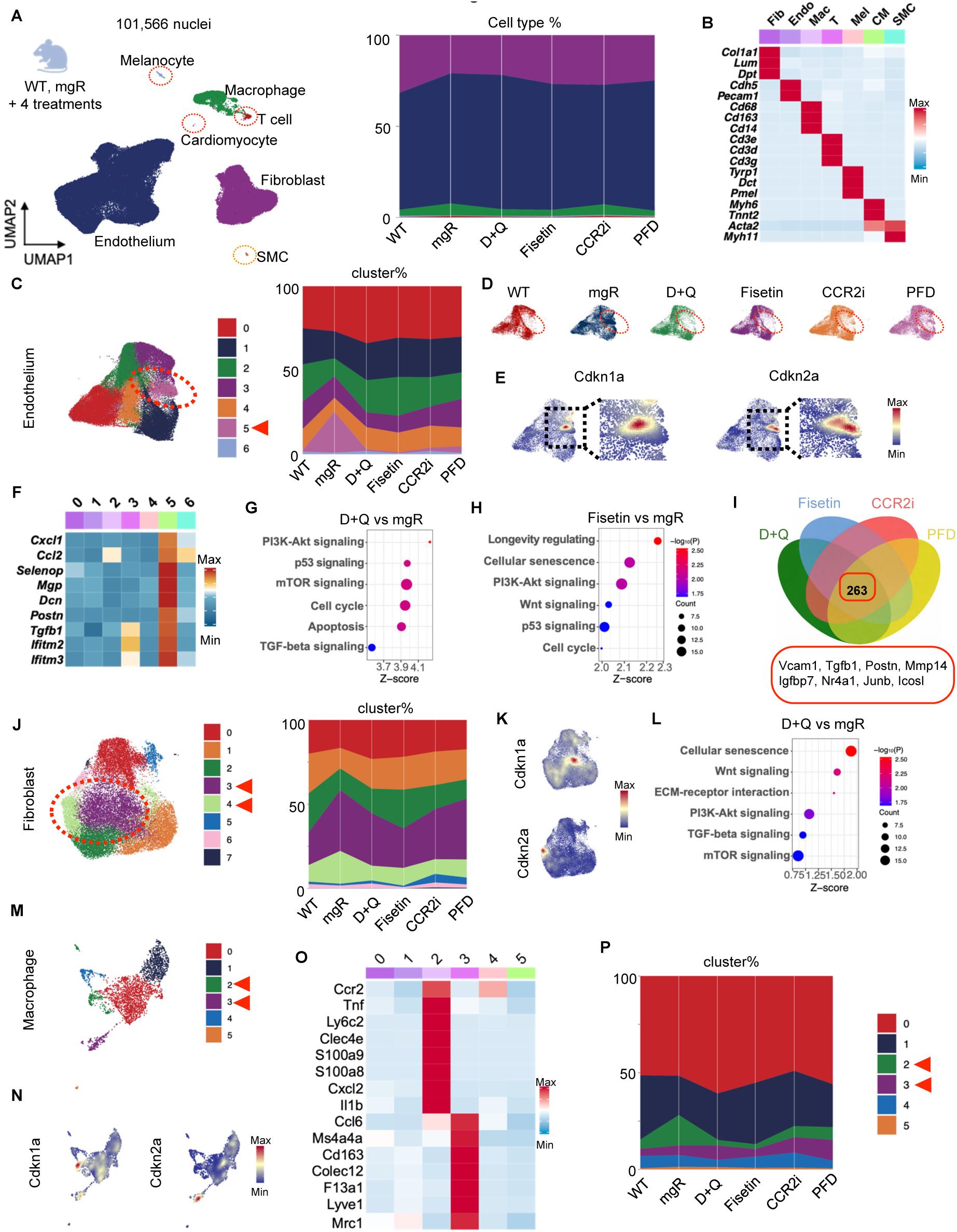
snRNA-seq reveals suppression of senescent cell populations following senolytic therapy in mitral valve disease. **(A)** UMAP showing the cellular composition of mitral valves across experimental groups (left) and stacked bar plots showing the proportional distribution of major cell types (right). **(B)** Expression of canonical marker genes across major cell types. Smooth muscle cells (SMC). **(C)** Unsupervised clustering of endothelial cells (left), with the circled cluster indicating a senescence-associated endothelial population characterized by enriched Cdkn1a and Cdkn2a expression; stacked bar plots show the proportional distribution of endothelial clusters across experimental groups (right). **(D)** UMAP showing abundance changes of senescence-associated endothelial cells across experimental groups. **(E)** Expression of Cdkn1a and Cdkn2a in endothelial cells, with magnified views of selected regions. **(F)** Heatmap of representative fibroinflammatory genes across endothelial clusters. **(G-H)** Bubble plots showing representative enriched KEGG pathways derived from DEGs in D+Q versus mgR and fisetin versus mgR comparisons. **(I)** Venn diagram showing the overlap of DEGs from D+Q versus mgR, fisetin versus mgR, CCR2i versus mgR, and PFD versus mgR comparisons, with 263 genes shared across all treatment groups. **(J)** Unsupervised clustering of fibroblasts (left), with circled clusters indicating a senescence-associated fibroblast population characterized by enriched Cdkn1a and Cdkn2a expression; stacked bar plots show the proportional distribution of fibroblast clusters across experimental groups (right). **(K)** Expression of Cdkn1a and Cdkn2a in fibroblasts. **(L)** Bubble plots showing representative enriched KEGG pathways derived from DEGs in D+Q versus mgR. **(M)** Unsupervised clustering of macrophages. **(N)** Expression of Cdkn1a and Cdkn2a in macrophages. **(O)** Heatmap showing expression of representative M1-like and M2-like macrophage-associated genes across macrophage clusters. **(P)** Stacked bar plots showing the proportional distribution of macrophage clusters across experimental groups. Data are presented as mean ± SEM. Statistical significance was determined using appropriate tests as described in the text.

In endothelial cells, unsupervised clustering identified seven transcriptionally distinct subpopulations (Figure 6C and Figure S7B). Endothelial cluster 5 was expanded in mgR valves, whereas only a small fraction of endothelial cells from WT valves occupied this state. This cluster almost disappeared in senolytic-treated valves (D+Q and fisetin) and was also reduced following CCR2i or pirfenidone treatment, albeit to a lesser extent (Figure 6C and 6D). Feature plots demonstrated enrichment of both *Cdkn1a* and *Cdkn2a* in cluster 5, indicating an endothelial population enriched for senescence-associated transcriptional features^80–82^ (Figure 6E). Notably, higher-resolution visualization revealed that *Cdkn1a*- and *Cdkn2a*-expressing endothelial cells were not identical within this cluster, again, suggesting distinct and coordinated *Cdkn1a*-and *Cdkn2a* senescent subsets. Concordantly, this endothelial cell cluster exhibited coordinated upregulation of fibroinflammatory and ECM-remodeling programs, consistent with a role in promoting immune recruitment and paracrine activation of fibroblasts (Figure 6F and Figure S7C). In fibroblasts, unsupervised clustering identified eight subpopulations (Figure 6J and Figure S8A). Two fibroblast clusters (clusters 3 and 4) were expanded in mgR valves and were reduced following senolytic treatment (Figure 6J). Feature plots showed that cluster 3 was enriched for *Cdkn1a*, whereas cluster 4 preferentially expressed *Cdkn2a* (Figure 6K). These two fibroblast clusters were enriched for inflammatory genes and exhibited upregulation of ECM-remodeling and profibrotic markers (Figure S8B). Senolytic therapies reduced both fibroblast clusters, whereas non-senolytic interventions preferentially reduced the *Cdkn2a*-high population, indicating state-specific therapeutic sensitivity among senescence-associated fibroblast subtypes. Although p16 (*Cdkn2a*) has been implicated in more severe and pathogenic senescent cell state, we show that it is required to target both *Cdkn1a and Cdkn2a* programs to achieve therapeutic efficacy.. Consistent with these cellular changes, pathway enrichment analyses demonstrated broad suppression of pathways commonly associated with senescence and senescence-induced functional alterations in senolytic-treated valves, including PI3K-AKT signaling, mTOR signaling, p53 pathways, cell-cycle regulation, and apoptosis (Figure 6G, 6H, and 6L). In addition to immune-related signaling and fibrotic remodeling pathways, CCR2i and pirfenidone also modulated senescence-associated pathways (Figure S7D, S7E, and S8C-S8E) but could not achieve the appreciable level of therapeutic efficacy (Figure 5). Cross-condition analysis identified shared transcriptional signatures across treatment groups. In endothelial cells, 263 genes, and in fibroblasts, 79 genes, were commonly regulated across interventions (Figure 6I, Figure S8F, and Tables S11 and S12) and were associated with ECM organization and fibroinflammatory remodeling programs.

In macrophages, clustering identified six subpopulations (Figure 6M). Feature plots showed that macrophage cluster 2 was enriched for *Cdkn1a*, whereas cluster 3 preferentially expressed *Cdkn2a* (Figure 6N). Marker gene analyses indicated that cluster 3 corresponded to an M2-like resident macrophage program, whereas cluster 2 represented a Ccr2⁺ M1-like monocyte-derived macrophage population^14,60–62^ (Figure 6O). Notably, both *Cdkn1a*- and *Cdkn2a*-associated macrophage populations were expanded in mgR valves, with *Cdkn1a*-associated Ccr2⁺ monocyte-derived macrophages showing a more pronounced increase than *Cdkn2a*-associated resident macrophages (Figure 6P). Following senolytic treatment, the *Cdkn1a*-associated macrophage population was markedly reduced, whereas CCR2 inhibition and pirfenidone resulted in modest reductions, although levels remained lower than in untreated mgR mice (Figure 6P). In contrast, *Cdkn2a*-associated macrophage populations were not consistently reduced across treatments and, under certain conditions, exhibited a modest increase in relative abundance. These findings are consistent with prior studies demonstrating that Ccr2^+^ monocyte-derived macrophages play a critical role in mitral valve degeneration and that targeting this population can attenuate disease progression.

To define how therapeutic interventions reshape senescence-associated cell-cell communication and intercellular signaling, we performed ligand-receptor interaction analyses. Comparative analyses between mgR valves and their corresponding treated counterparts revealed a common attenuation of intercellular communication networks. Downregulated interactions spanned multiple cell-cell axes and converged on TGF-β signaling and ECM-integrin interactions, which are known to be associated with fibrosis, as well as immune regulatory pathways, consistent with coordinated suppression of inflammatory and fibrotic remodeling signals across cell types (Figure S9 and S10).

### Conserved senescence programs involving both *CDKN1A and CDKN2A* in other ECM-dominant human diseases

We define ECM-dominant cardiovascular diseases as conditions in which remodeling and disruption of ECM architecture including fibrosis and degeneration is a primary determinant of structural failure. To assess whether senescence-associated programs converge across this disease class, we analyzed three representative ECM-dominant cardiovascular pathologies using public datasets: calcific aortic valve disease (CAVD)^83^ and thoracic aortic aneurysm (TAA) in both human (hTAA)^84^ and mouse (mTAA)^85^ models, as well as our mouse whole heart cell mapping and senotypic treatment study.

We first examined CAVD, in which four major cell types were identified based on established marker genes (Figure S11A and S11B). Given the central role of fibroblasts in valvular ECM remodeling, subsequent analyses focused on this population. Unsupervised clustering resolved nine distinct fibroblast clusters (Figure 7A). Unsupervised cluster-enriched top marker genes defined distinct transcriptional states, with cluster 0 characterized by an inflammatory program and cluster 3 by an ECM-remodeling, profibrotic program (Figure 7B). Consistently, *CDKN1A* was preferentially enriched in cluster 0, whereas *CDKN2A* was enriched in cluster 3 (Figure 7C). In line with these patterns, the *CDKN1A*-associated cluster 0 showed increased expression of inflammatory genes, including *CCL2, IL6*, and multiple *CXCL* chemokines, whereas the *CDKN2A*-associated cluster 3 expressed higher levels of profibrotic and ECM-related genes such as *COL3A1, POSTN*, and *FN1* (Figure 7D).

**Figure 7.**
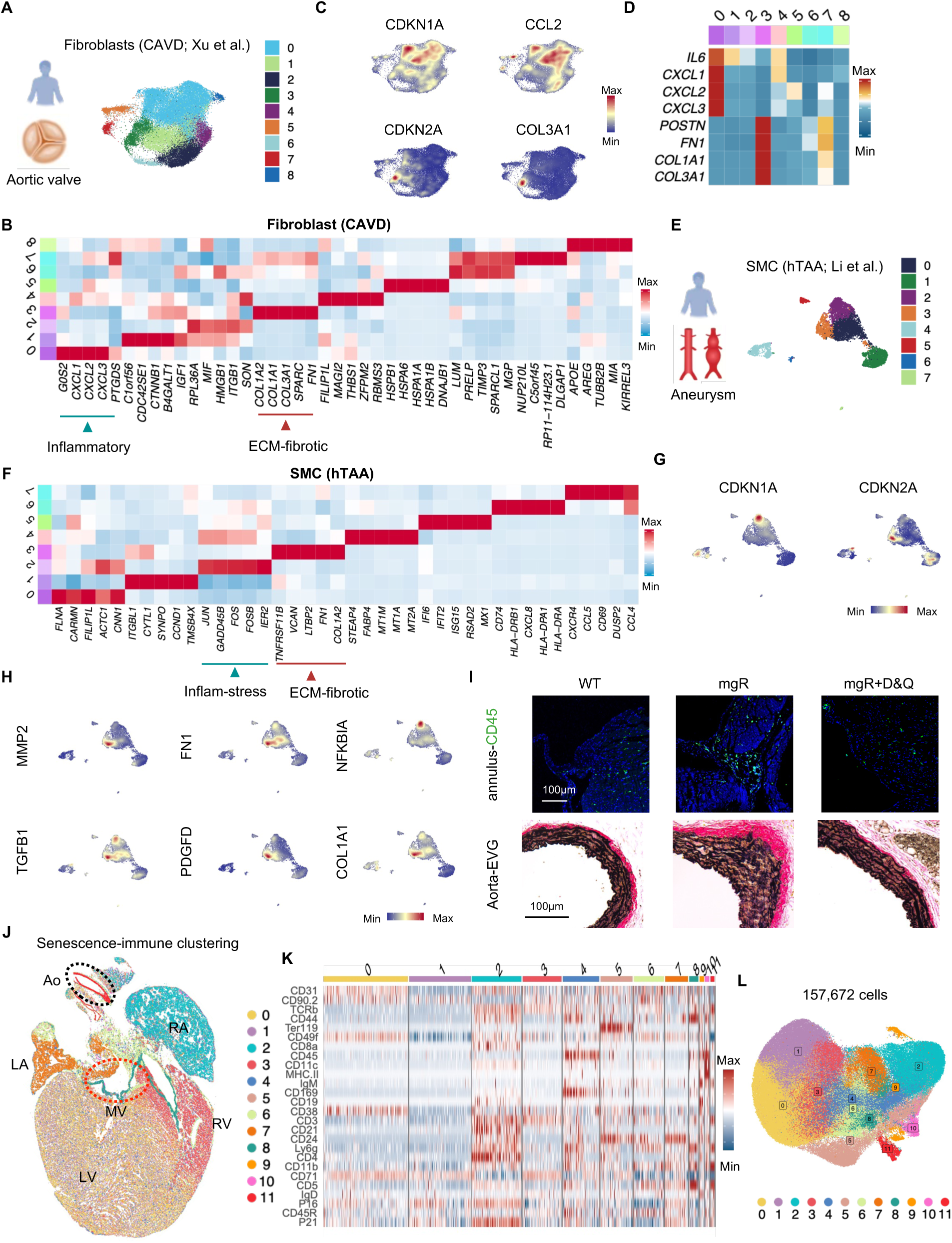
Conserved senescence programs underlie ECM-dominant cardiovascular diseases and are targetable by senolytic therapies. **(A)** Unsupervised clustering of fibroblasts from calcific aortic valve disease (CAVD). **(B)** Heatmap showing the top five marker genes for each fibroblast cluster. **(C)** Expression of CDKN1A, CDKN2A, CCL2, and COL3A1 across fibroblasts from CAVD. **(D)** Heatmap of representative inflammatory and fibrotic ECM-remodeling genes across fibroblast clusters. **(E)** Unsupervised clustering of SMCs from human thoracic aortic aneurysm (hTAA). **(F)** Heatmap showing the top five marker genes for each SMC cluster. **(G)** Expression of CDKN1A and CDKN2A across SMCs from hTAA. **(H)** Expression of FN1, TGFB1, MMP2, COL1A1, NFKBIA, and PDGFD across SMCs from hTAA. **(I)** Representative immunofluorescence staining of CD45 in the valve annulus (top), and representative histological features of the thoracic aorta across the indicated experimental groups (bottom). **(J)** Unsupervised clustering of mouse whole-heart tissue based on CODEX imaging of senescence and immune markers. **(K)** Heatmap showing the normalized expression of marker proteins across clusters. **(L)** UMAP visualization of unsupervised clustering of 157,672 cells profiled by CODEX imaging. Data are presented as mean ± SEM. Statistical significance and methods are described in the main text.

We next analyzed human TAA, in which nine major cell types were identified (Figure S11C and S11D). Given the critical role of smooth muscle cells (SMCs) in maintaining aortic wall integrity and ECM homeostasis, analyses focused on this population. Unsupervised clustering resolved eight distinct SMC clusters (Figure 7E and 7F). Unsupervised cluster-enriched top marker genes indicated that the cluster 2 exhibited an inflammatory and stress-responsive program, whereas the cluster 3 showed a fibrotic and ECM-oriented profile (Figure 7F). Consistent with these programs, *CDKN1A* was preferentially enriched in cluster 2, whereas *CDKN2A* was enriched in cluster 3 (Figure 7G). Consistent with their functional identities, these senescence-associated clusters were enriched for key pathogenic mediators of TAA, including NF-κB-associated inflammatory activation (*NFKBIA*), growth factor-driven signaling (*TGFB1, PDGFD*), and ECM remodeling coupled with proteolytic turnover (*FN1, COL1A1, MMP2*) (Figure 7H)

To assess cross-species conservation, we also examined mouse TAA. Four major cell types were identified (Figure S11E and S11F), and analyses again focused on SMCs. Unsupervised clustering resolved seven SMC clusters (Figure S11G). Mirroring human TAA, *Cdkn1a* was preferentially enriched in cluster 0, whereas *Cdkn2a* was enriched in cluster 2 (Figure S11H). Moreover, senescence-associated SMC clusters similarly exhibited enrichment of key pathogenic mediators (Figure S11H).

We next assessed the impact of senolytic therapy across cardiovascular tissues beyond the mitral valve in the mgR mouse model. mgR mice were treated with D+Q for 8 weeks (4-12 weeks of age). Senolytic treatment ameliorated aortic pathology, improving elastic lamellar organization and reducing medial degeneration compared with untreated mgR mice (Figure 7I). In parallel, D+Q reduced CD45⁺ immune cell accumulation in the valve annulus and fibrotic cardiac skeleton, regions that exhibited prominent immune infiltration in untreated mgR mice (Figure 7I).

Although senescence is a major risk factor across cardiovascular diseases, senescence-associated features vary intrinsically across tissues and are not uniformly distributed across anatomical locations or tissue contexts. To define their spatial organization, we performed CODEX multiplex imaging on whole mouse heart using a combined senescence and immune antibody panel. Spatial clustering analysis revealed that distinct cardiac regions, including the four chambers, valves, fibrotic cardiac skeleton, and aorta, segregated into discrete senescence- and immune-associated microenvironments (Figure 7J–7L). Together, these findings further support that senescence-associated programs are deployed to drive a range of cardiovascular pathologies only in spatially specialized regions, shaped by regional structural demands and local ECM context.

## DISCUSSION

In this study, we identify cellular senescence as a central pathogenic driver in ECM-dominant cardiovascular pathologies and delineate how senescence-associated programs coordinate inflammatory signaling with maladaptive ECM remodeling. By integrating human and murine single cell transcriptomics with spatial mapping and interventional analyses, we demonstrate that the vascular fibrotic disease is not merely a passive consequence of age-associated ECM damage accumulation but an actively maintained pathological state sustained by at least two distinct and coordinated senescence-associated cellular programs. Notably, selective elimination of senescent cells more effectively restores valve structure and function than pathway-specific anti-inflammatory or antifibrotic strategies, establishing senescence as a therapeutically actionable upstream driver of degenerative valvular disease.

A key conceptual advance of this work is the resolution of distinct senescence programs and senescence heterogeneity within fibroblasts. We identify two dominant, non-redundant senescence-associated programs marked by *CDKN1A* or *CDKN2A*. These programs do not represent interchangeable stages of a uniform endpoint but instead define functionally specialized states with distinct effector outputs. *CDKN1A*-associated fibroblasts are enriched for inflammatory and stress-responsive pathways, including chemokine production and immune recruitment, whereas *CDKN2A*-associated fibroblasts preferentially exhibit profibrotic ECM remodeling and matrix deposition programs. Through paracrine and juxtacrine signaling, these states reinforce each other and shape the surrounding microenvironment, forming a feed-forward circuit that couples immune activation to progressive ECM disorganization^1,3,5,86,87^.

Senescence-associated reprogramming extends beyond fibroblasts to the immune compartment, establishing a coordinated fibroinflammatory network. The diseased tissue is characterized by depletion of cytotoxic CD8⁺ T cells and expansion of inflammatory and chronically activated T cell populations, including *STAT4*⁺ CD4⁺ T cells and *GZMK*⁺ effector-memory CD8⁺ T cells. In parallel, macrophages adopt distinct senescence-associated states, including *CCR2*⁺ *CDKN1A*-associated monocyte-derived macrophages and *CDKN2A*-associated resident macrophages, consistent with differential roles in inflammatory amplification and matrix remodeling. These findings support a model in which immune cell reprogramming is tightly coupled to fibroblast senescence, collectively sustaining or amplifying chronic inflammation and tissue remodeling.

These immune alterations are accompanied by increased expression of immune checkpoint ligands and anti-apoptotic pathways in fibroblasts, consistent with impaired immune-mediated clearance^32,33,40^. Such reprogramming promotes persistence of senescence-associated cells through resistance to cytotoxic T cell activity and reduced macrophage-mediated clearance^32,33,36,40^. Together, these observations indicate that senescent fibroblast states are not passive endpoints but actively maintain and remodel their microenvironment.

Collectively, our data further support a model in which senescence is deployed across multiple interacting cell types rather than confined to a single “driver” population while senescent fibroblasts rather than myofibroblasts serve as the origin of this cascade. In this framework, senescence functions as a microenvironmental organizer that establishes and stabilizes a permissive niche for sustained fibroinflammatory remodeling.

Mechanistically, ligand-receptor analyses and spatial mapping further indicate the existence of an organized senescence-associated communication network, with fibroblasts acting as a central signaling hub. These interactions involve pathways governing ECM remodeling, survival, and immune regulation, and are spatially structured such that senescence markers, immune infiltrates, and profibrotic remodeling co-localize within defined microenvironments. In this context, integrin-mediated mechanotransduction and downstream signaling pathways, including TGF-β, likely serve as key intermediates linking ECM disorganization to stabilization of senescence programs. These findings define a mechanically coupled senescence network that integrates inflammatory and fibrotic processes into a unified pathogenic continuum.

Although *CDKN1A* is frequently associated with early or stress-induced senescence and *CDKN2A* with more stable remodeling-associated programs, our data do not directly establish a lineage relationship between *CDKN1A*-high and *CDKN2A*-high fibroblast states. Accordingly, we cannot determine whether *CDKN2A*-high fibroblasts arise from *CDKN1A*-high precursors or represent independent senescent trajectories, which requires future studies such as lineage-tracing experiments. In contrast, within the immune compartment, *CDKN1A*- and *CDKN2A*-associated programs align with distinct differentiation and polarization states. Nevertheless, our study highlights the need for both *CDKN1A*-high and *CDKN2A*-high populations to co-exist either as distinct lineages or dynamic states to coordinate and drive pathogenesis.

Our findings further indicate that senescence in valvular heart disease is not solely driven by chronological aging. While senescent cell accumulation correlates with age in human tissues, robust senescence programs observed in young mgR mice indicate that pathological stress can induce premature senescence. In this setting, fibrillin-1 deficiency likely generates sustained biomechanical stress and dysregulated TGF-β signaling, creating a chronic stress niche that promotes senescence and fibroinflammatory remodeling^70^. The convergence of these programs across species supports senescence as a common pathological endpoint arising from diverse upstream insults.

Cellular senescence has traditionally been categorized into replicative, oncogene-induced, and stress-induced forms. Consistent with this, the senescence landscape in our study does not conform to a single canonical category but instead reflects a composite stress-associated program integrating mechanical strain, ECM disorganization, inflammatory signaling, and metabolic stress^88,89^.

The mgR model further enabled direct comparison of therapeutic strategies targeting senescence, inflammation, and fibrosis. While CCR2 inhibition and pirfenidone attenuated immune recruitment and ECM accumulation, they incompletely suppressed senescence-associated states and produced limited structural recovery and minimal physiological function restoration. In contrast, senolytic therapies (D+Q or fisetin) robustly reduced both *CDKN1A*+ and *CDKN2A+* senescent cell burden, restored normal ECM organization, decreased inflammatory infiltration, and improved valve function, including appreciable benefit even when administered after disease onset. These findings support a model in which senescence acts upstream of inflammatory and fibrotic remodeling, and suggest that the conventional pathway-restricted interventions are insufficient to durably prevent or reverse the disease trajectory.

Importantly, therapeutic effects were cell type- and program-specific. Senolytics reduced both *CDKN1A*- and *CDKN2A*-associated fibroblast populations, whereas non-senolytic interventions preferentially affected *CDKN2A*-enriched states. Interesting, in macrophages, senolytics selectively reduced *CDKN1A*-associated CCR2⁺ populations, with some effects on *CDKN2A*-associated resident macrophages. These findings highlight functional heterogeneity within senescence-associated programs and suggest opportunities for more precise senolytic drugs for therapeutic targeting.

Beyond valvular heart disease MMVD, conserved senescence-associated *CDKN1A* and *CDKN2A* programs were identified across multiple ECM-dominant cardiovascular diseases, including CAVD and TAA, supporting a generalizable fibroinflammatory remodeling mechanism^90–92^. Importantly, spatial mapping indicates that these programs are deployed in a region-specific manner across the heart and great vessels, shaped by local ECM context and immune microenvironments. Such spatial specialization may help explain the tissue-selective manifestations of senescence-associated degeneration and may further account for why pathway-specific cytokine blockade, such as anti-IL1B therapy, can be effective in certain cardiovascular contexts, including heart failure, yet shows limited benefit in others, such as MMVD^78^.

Several limitations should be acknowledged. First, valvular heart diseases progress over decades in humans but over weeks in mice, which may influence the dynamics and composition of senescence-associated programs. Second, due to the dense ECM of the mitral valve and the use of a mild enzymatic dissociation protocol (10-15 min), nuclei isolation might result in skewed cell-type representation. Cell types that are more readily dissociated are disproportionately enriched, whereas cells more deeply embedded within the ECM and resistant to dissociation are relatively underrepresented. This dissociation-related bias primarily affects estimates of cell-type proportions; however, it is unlikely to alter cell-intrinsic transcriptional programs or confound disease-associated state comparisons within individual cell types. Third, although senolytic therapies were effective in experimental models, the durability of benefit, optimal dosing regimens, and long-term safety remain to be established in humans. Finally, while we quantified senescence heterogeneity characterized by *CDKN1A*- and *CDKN2A*-associated programs, both their upstream regulatory drivers and their gene-specific functional contributions remain incompletely resolved. Dissecting the causal roles of these programs will require temporal perturbation strategies, including targeted genetic modulation and the use of *CDKN1A*- and *CDKN2A*-specific genetically modified models.

In addition, although our analyses define a coordinated senescence-ECM degeneration axis at single-cell resolution, this study was designed to resolve organizational and cellular-state architecture rather than to dissect specific signaling pathways governing program initiation or intercellular hierarchy. The relative contributions of individual pathways to the establishment and maintenance of *CDKN1A*- and *CDKN2A*-associated states remain to be determined through targeted genetic and mechanistic perturbation studies. Future work integrating pathway-specific modulation with lineage tracing and temporal analyses will be required to define causal relationships within the senescence circuit.

In summary, our findings redefine ECM-dominant cardiovascular pathologies as a senescence-driven disease characterized by coordinated deployment of distinct *CDKN1A*- and *CDKN2A* senescence programs across multiple cell types. Rather than a passive consequence of aging, senescence emerges as an active organizing force that integrates inflammatory signaling with ECM remodeling in disease progression. The ability of senolytic therapies to reverse structural and functional pathology establishes a direct causal link between senescence and disease maintenance, providing a rationale for targeting senescence as a unified therapeutic strategy in these ECM-dominant cardiovascular disorders.

## METHODS

### Ethics

Animal studies were approved by the Institutional Animal Care and Use Committee (IACUC) of Yale University. Mice were housed under standardized conditions in the Yale University Animal Facility, and all experimental procedures were performed in accordance with applicable U.S. federal regulations. Human tissue collection was approved by the Institutional Review Board (IRB) of Yale University and conducted in compliance with U.S. regulations governing the ethical use of human tissues.

### Mice

Fbn1^mgR^/^mgR^ mice^1^ (stock no. 005704; The Jackson Laboratory) were used in this study and are referred to as mgR mice. All mouse strains were maintained on a C57BL/6J genetic background. Mice were euthanized at the indicated ages for downstream analyses. Wild-type (WT) control hearts were collected from sex-matched littermates within each experimental group. Both male and female mice were used in this study unless otherwise indicated, and groups were sex-matched within experiments. For senolytic treatment, dasatinib (Cayman Chemical; 5 mg/kg) and quercetin (MedChemExpress; 50 mg/kg), or vehicle control, were administered by oral gavage for three consecutive days every two weeks^2^. For fisetin treatment, fisetin (STEMCELL Technologies; 100 mg/kg) or vehicle control was administered by oral gavage for three consecutive days every two weeks^3,4^. This intermittent dosing strategy was chosen based on prior studies demonstrating effective senescent cell clearance while minimizing off-target effects. The CCR2 antagonist RS504393 (MedChemExpress; 2 mg/kg) or vehicle control was administered by daily intraperitoneal injection^5^. For IL-1β neutralization, mice were treated with either an isotype control antibody (Bio X Cell, BE0091) or an anti-IL-1β neutralizing antibody (Bio X Cell, BE0246) at a dose of 5 mg/kg by intraperitoneal injection every three days^6,7^. Pirfenidone (MedChemExpress; 300 mg/kg) or vehicle control was administered by oral gavage once daily as previously described^8^. Animals were randomly assigned to treatment groups, and investigators were blinded to treatment allocation during data acquisition and analysis whenever feasible. Treatments were initiated at the indicated ages and continued until tissue harvest, as specified for each experimental cohort.

### Echocardiography

Mitral valve function was assessed by high-resolution transthoracic echocardiography in animals anesthetized with light isoflurane. A 40-MHz linear array transducer (Vevo 2100 system, VisualSonics) was used to detect mitral regurgitation. Echocardiography was performed by an investigator blinded to genotype and treatment.

### Gross morphology assessment

Mouse hearts were excised and fixed in 4% paraformaldehyde at 4 °C overnight. Following fixation, the mitral valves were carefully exposed for gross morphological assessment. Quantitative morphometric analyses were performed using ImageJ software.

### Histomorphometry

Tissues were fixed in 4% paraformaldehyde overnight, paraffin-embedded, and sectioned at a thickness of 7 μm. Histological staining of paraffin sections was performed by the Yale Research Histology Laboratory using standard protocols, including Hematoxylin and Eosin (H&E), Elastin van Gieson (EVG), Masson’s trichrome, and Movat’s pentachrome staining. Senescence-associated β-galactosidase staining was performed on cryosections using standard protocols. Quantitative morphometric analyses were performed using ImageJ software.

### Confocal imaging

Tissues were embedded in OCT and sectioned at 7 μm. Sections were washed with Tris-buffered saline (TBS) and incubated overnight at 4 °C with primary antibodies diluted in blocking solution (10% BSA and horse serum in TBS). After TBS washes, sections were incubated with Alexa Fluor 488-, 594-, or 647-conjugated secondary antibodies for 1 hour at room temperature, washed, and mounted with ProLong Gold Antifade Mountant with DAPI (Thermo Fisher Scientific, P36935). Primary antibodies used in this study included anti-CD45 (R&D, AF114), anti-CCL2 (Abcam, ab25124), anti-p16 (Cell Signaling Technology, 88667, 23200), anti-p21 (Cell Signaling Technology, 2947, 39256), and anti-collagen III (Abcam, ab7778). Primary antibodies were used at manufacturer-recommended dilutions. Images were acquired using a Leica SP8 confocal microscope. ImageJ software was used to quantify immunofluorescence signals by calculating the mean fluorescence intensity, defined as the sum of pixel values within each valve section normalized to the measured area and expressed as arbitrary units (AU) or number of positive stained cells for specific antibodies.

### RNA-seq

Total RNA was isolated from mitral valve tissues and used for library preparation with the RNA Library Prep Kit for Illumina (NEB, E6420L), with cDNA synthesis performed according to the manufacturer’s protocol. Libraries were constructed using the NEBNext Ultra II workflow, including fragmentation, end repair, adapter ligation, and PCR amplification. Indexed libraries passing quality and quantity thresholds were quantified by qPCR (KAPA Biosystems), and libraries were sequenced on a NovaSeq 6000 system (Illumina) at the Yale Center for Genome Analysis. Sequencing reads were aligned to the mouse (mm10) or human (hg38) reference genome, and read counts were normalized by the trimmed mean of M-values (TMM) method with differential expression analysis performed using edgeR.

### snRNA-seq and raw data processing

Nuclei were isolated from mitral valve tissue using the Chromium Nuclei Isolation Kit (10x Genomics; PN-1000493), stained with 7-AAD, FACS-sorted, and collected in PBS containing 0.4% BSA. snRNA-seq libraries were generated using the Chromium Single Cell Gene Expression platform (10x Genomics) according to the manufacturer’s protocol. Due to the dense extracellular matrix of mitral valves and the use of a brief enzymatic dissociation protocol (10-15 min), nuclei isolation introduced biases in cell-type representation. In murine samples, the small size of the mitral valve precluded tissue subdivision, necessitating dissociation of whole valves. This resulted in relative enrichment of endothelial nuclei, consistent with their surface localization and greater susceptibility to dissociation. In human samples, although valve tissue could be subdivided prior to processing, immune cells were relatively enriched, likely reflecting their higher ease of liberation during dissociation. In contrast, fibroblasts were comparatively underrepresented across datasets, consistent with their strong embedding within the ECM and relative resistance to dissociation. Importantly, this dissociation-associated bias is expected to affect estimates of cell-type proportions but is unlikely to alter cell-intrinsic transcriptional programs or confound disease-associated state comparisons performed within individual cell types. Raw data were processed with Cell Ranger^9^.

### snRNA-seq data analysis

Analyses were performed in R using Seurat (v4.3.0). Nuclei with fewer than 200 or more than 5,000 detected genes, or with mitochondrial gene expression exceeding 10%, were excluded^10,11^. Expression data were normalized using LogNormalize, dimensionality reduction was performed by principal component analysis on highly variable genes, and batch correction was performed using Harmony (v1.1.0). UMAP was used for visualization and differentially expressed genes were defined by log fold change >0.25 and adjusted P < 0.05. Gene expression matrices were extracted from RDS files and ligand-receptor interactions were inferred using CellChat^12^. Pathway enrichment analysis was performed on differentially expressed genes using clusterProfiler, and pathways with an adjusted P value < 0.05 were considered statistically significant. For pathway analyses, enrichment was summarized using Z-scores derived from normalized enrichment statistics, while statistical significance was represented as -log10(adjusted P value). Cell-type and cluster proportions were compared using samples as the unit of replication with propeller (speckle) on arcsine square-root-transformed proportions, with Benjamini-Hochberg FDR correction. To quantify distinct senescence-associated functional programs, we derived gene-module scores representing inflammatory-stress signaling, pro-fibrotic/extracellular matrix remodeling, and senescent survival. Gene sets were anchored on the SenMayo senescence signature and extended with additional genes curated from prior literature to capture valve-relevant and cell-type-specific inflammatory, survival, and extracellular matrix remodeling programs (Table S10)^13^. Gene-module scores were calculated at the single-cell level using the AddModuleScore function in Seurat. Scores were calculated using identical parameters across datasets and compared within annotated cell types.

### Spatial-CITE-seq

100×100 20um chip was used for human mitral valve spatial CITE-seq profiling, as previously described^14–16^. Spatial omics libraries were constructed and sequenced on an Illumina NovaSeq 6000 system. For cDNA derived from mRNA, raw FASTQ files containing unique molecular identifiers (UMIs) and spatial barcodes (barcode A and barcode B) were reformatted into the input structure required by the ST Pipeline (v1.7.2) using a custom Python script^17^. Using recommended ST Pipeline parameters, RNA and protein expression matrices were generated and subsequently clustered in Seurat^18^. Transcriptomic data were normalized using the SCTransform function in Seurat, whereas protein expression data were normalized using the centered log-ratio (CLR) transformation method.

### Multiplexed imaging by CO-Detection by indexing (CODEX)

Tissues were embedded in OCT compound and sectioned at a thickness of 7 μm. Sections were stained with DNA-barcoded antibodies and subjected to iterative fluorophore cycling for multiplexed imaging. Images were acquired across multiple imaging cycles using a fluorescence microscope, with acquisition parameters optimized for each fluorophore as previously described^19^. CODEX data were processed using StarDist in QuPath for cell detection^20,21^. The resulting data were subsequently analyzed using Seurat, including normalization, scaling, and principal component analysis (PCA). Followed by unsupervised graph-based clustering with the FindClusters function.

### Quantitative RT-PCR

Total RNA was isolated from mitral valve tissues using the RNeasy Mini Kit (Qiagen, 74104). cDNA synthesis was performed using the High-Capacity cDNA Reverse Transcription Kit (ThermoFisher Scientific, 4368814), and quantitative PCR was carried out using TaqMan Gene Expression Master Mix (ThermoFisher Scientific, 4369016) according to the manufacturer’s instructions. The amplified genes and primer catalogs (ThermoFisher Scientific) were as follows: CDKN2A (Hs00923894_m1), CDKN1A (Hs00355782_m1), CCL2 (Hs00234140_m1), IL6 (Hs00174131_m1), TGFB1 (Hs00998133_m1), COL1A1 (Hs00164004_m1), COL3A1 (Hs00943809_m1), ACTB (Hs99999903_m1), Cdkn2a (Mm00494449_m1), Cdkn1a (Mm04205640_g1), Ccl2 (Mm00441242_m1), Col3a1 (Mm00802300_m1) and Actb (Mm02619580_g1). Quantitative PCR reactions were performed on a Bio-Rad CFX96 real-time PCR system. Gene expression levels were normalized to ACTB and calculated using the 2^-ΔΔCt^ method.

### Statistics and visualization

Data are presented as individual data points with lines indicating the mean ± standard error of the mean (SEM). Comparisons between two groups were performed using two-tailed student’s t-tests, and comparisons among multiple groups were conducted using one-way analysis of variance (ANOVA) followed by Tukey’s post hoc test when appropriate. A two-sided P value < 0.05 was considered statistically significant. All statistical analyses and graphical representations were generated using Prism version 10.0.0 (GraphPad Software). Box plots display the median, interquartile range (IQR), and minimum and maximum values, with whiskers extending to 1.5× IQR from the quartiles and outliers shown as individual points. Violin plots depict data distributions, with embedded box plots indicating the median and IQR; whiskers extend to 1.5× IQR, and outliers are shown as individual points.

### Illustrations

Schematic illustrations were created with BioRender.com and Adobe Illustrator.

### Publicly available scRNA-seq datasets

Publicly available scRNA-seq datasets were obtained from previously published studies. The mouse TAA dataset was from Pedroza et al. (2020)^22^. and the human TAA dataset from Li et al. (2020)^23^. A human CAVD dataset was obtained from Xu et al. (2020)^24^. Raw or processed data, as available, were downloaded from the original repositories and reanalyzed using a standardized preprocessing and downstream analytical pipeline to ensure consistency across datasets.

## Data and code availability

The datasets and custom code generated in this study are being prepared for public deposition and will be made available in a public repository prior to manuscript acceptance.

## Materials & Correspondence

All correspondence and request for materials should be addressed to R.F..

### ACKNOWLEDGEMENTS

We are grateful to all the participants in this study. This research was supported by the US National Institutes of Health grants (UG3CA257393, UH3CA257393, U54AG076043, U54AG079759) to R.F., the US National Institutes of Health grant (NIH R35 GM150838 and NIH R01 HL173271) to Y.L..

## AUTHOR CONTRIBUTIONS

Conceptualization, F.G., A.G., Y.L., and R.F.; experiments, F.G., A.F., D.Z., X.L., X.T., N.F., and M.Z.; data analysis, F.G., A.F., D.Z., X.L., J.N., M.Y., X.T., N.F., D.W., G.L., S.H., X.D., and M.Z.; original draft, F.G. and A.F.; review and editing, F.G., A.G., Y.L., and R.F..

