## Supplementary Figures for "Decoding and Targeting Coordinated CDKN1A and CDKN2A Senescence Programs in ECM-Dominant Cardiovascular Pathologies"

A

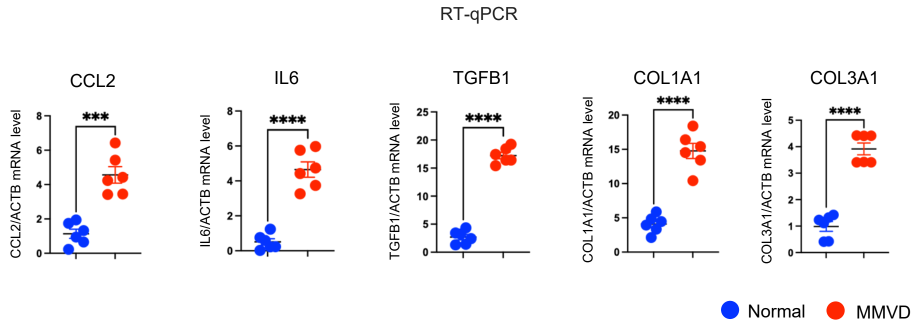


**Figure S1: Further analysis of myxomatous mitral valve disease**
**(A)** Scatter plots showing mRNA expression levels measured by RT-qPCR. Data are presented as mean ± SEM. ***p<0.001 and **** p<0.0001, by unpaired t-test.

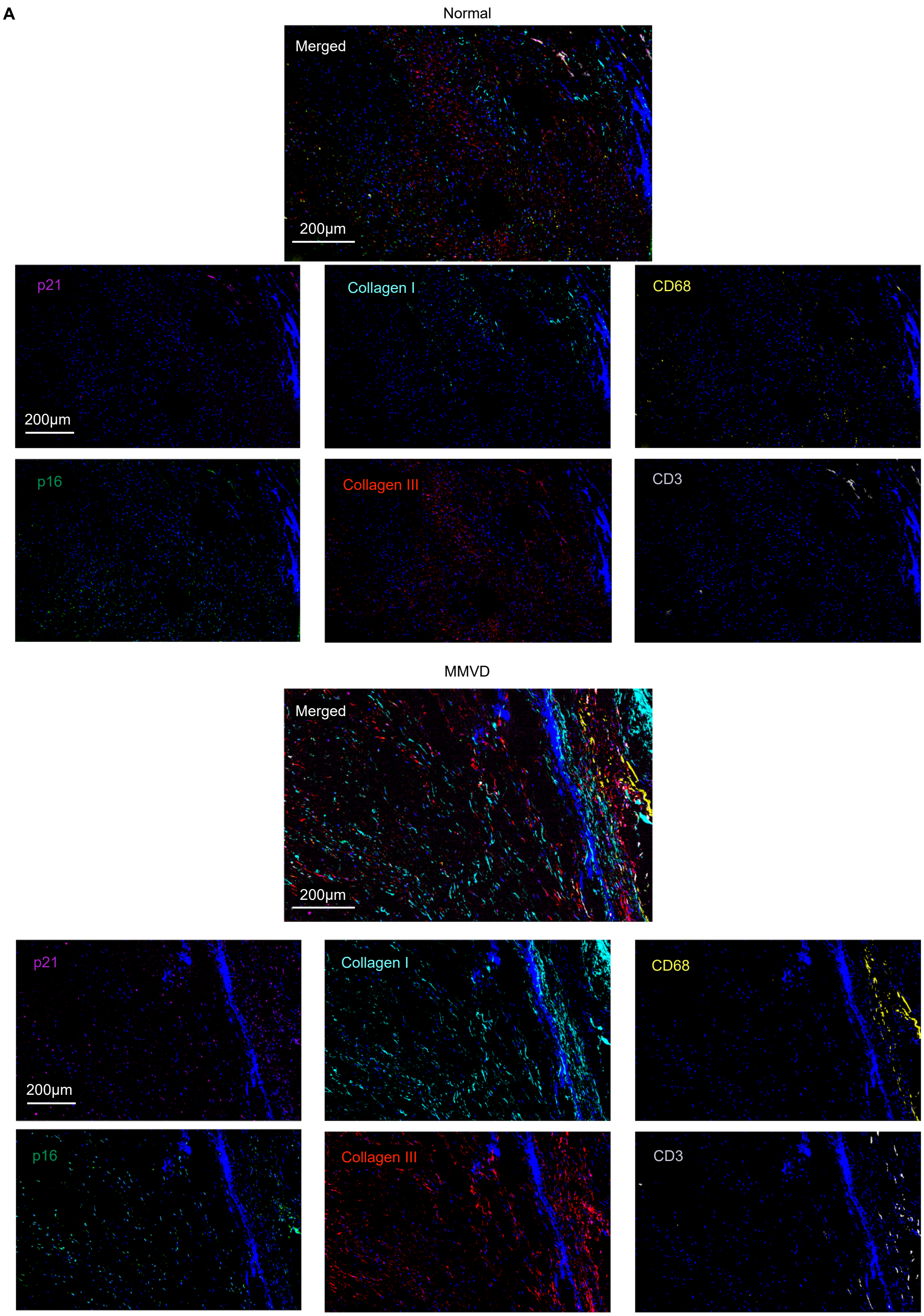


**Figure S2: Further characterization of myxomatous mitral valve disease**
**(A)** Representative CODEX multiplexed imaging of normal and myxomatous mitral valve tissues.

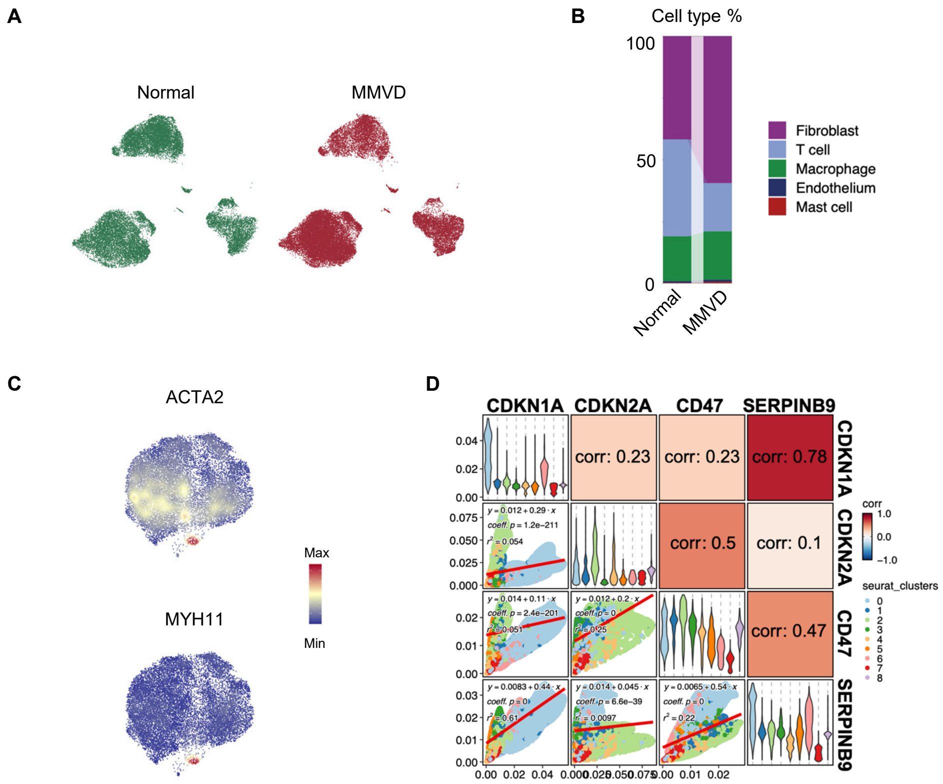


**Figure S3: Further characterization of myxomatous mitral valve disease**

**(A)** UMAP of all mitral valve cells from normal and MMVD samples.

**(B)** Stacked bar plots showing the proportional distribution of major cell types in normal and MMVD samples.

**(C)** Expression of canonical myofibroblast markers across fibroblast clusters.

**(D)** Correlation analysis of CDKN1A, CDKN2A, CD47, and SERPINB9 across fibroblast clusters.

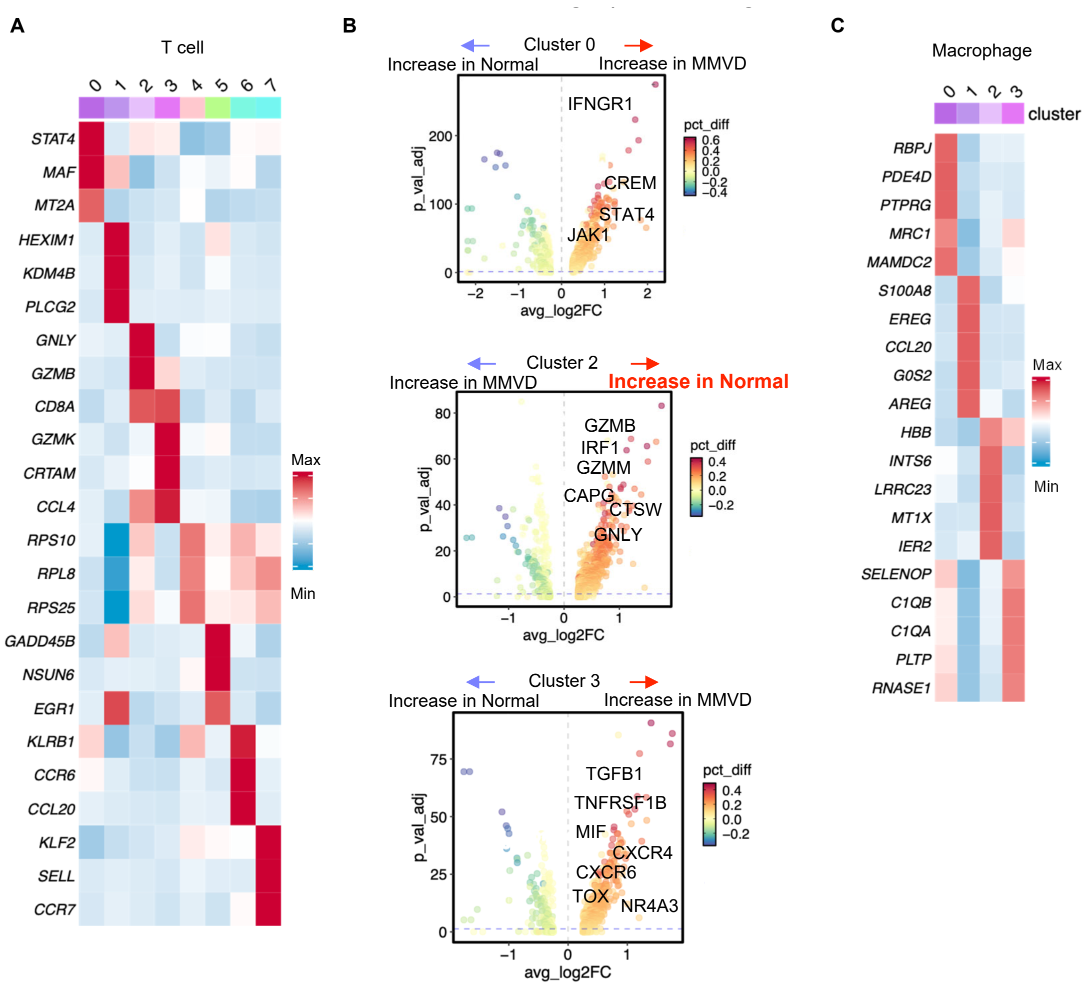


**Figure S4: Additional analyses of immune cells**

**(A)** Heatmap showing the top marker genes for each T cell cluster.

**(B)** Volcano plots showing cluster-specific DEGs between normal and MMVD T cells, with representative genes indicated.

**(C)** Heatmap showing the top marker genes for each macrophage cluster.

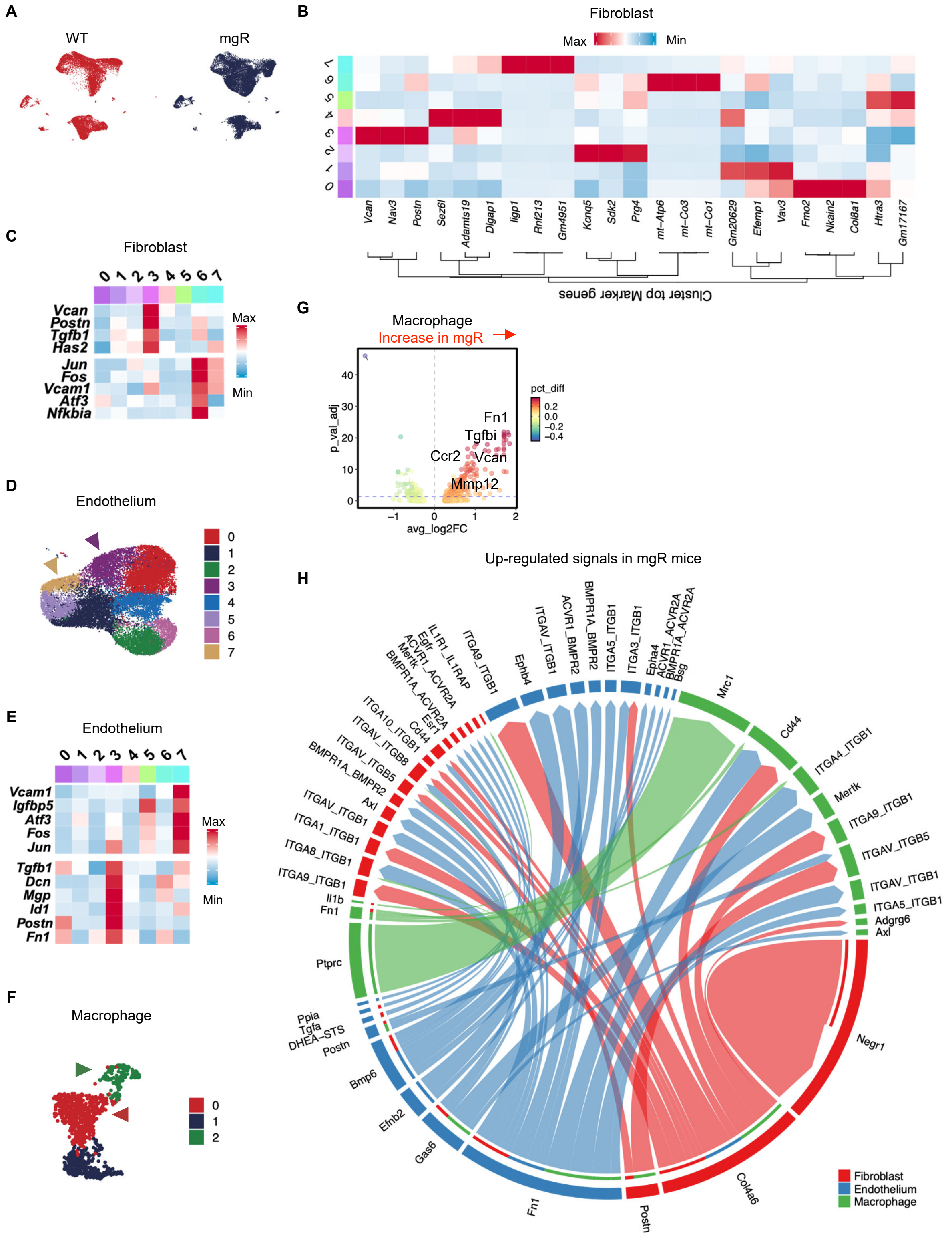


**Figure S5: Extended characterization of the mgR mouse model**

**(A)** UMAP of all mitral valve cells from WT and mgR.

**(B)** Heatmap showing the top marker genes for each fibroblast cluster.

**(C)** Heatmap showing representative marker genes for fibroblast cluster 3 (Cdkn2a) and 6 (Cdkn1a).
**(D)** Unsupervised clustering of endothelial cells. Arrows indicate endothelial clusters enriched for Cdkn1a and Cdkn2a expression.
**(E)** Heatmap showing representative fibroinflammatory marker genes for endothelial cell cluster 3 (Cdkn2a) and 7 (Cdkn1a).
**(F)** Unsupervised clustering of macrophages. Arrows indicate macrophage clusters enriched for Cdkn1a and Cdkn2a expression.
**(G)** DEGs between macrophages from WT and mgR mice. Representative genes upregulated in mgR macrophages are indicated.
**(H)** Representative ligand-receptor interactions upregulated in mgR compared with WT mouse mitral valves. Edge thickness reflects inferred interaction strength.

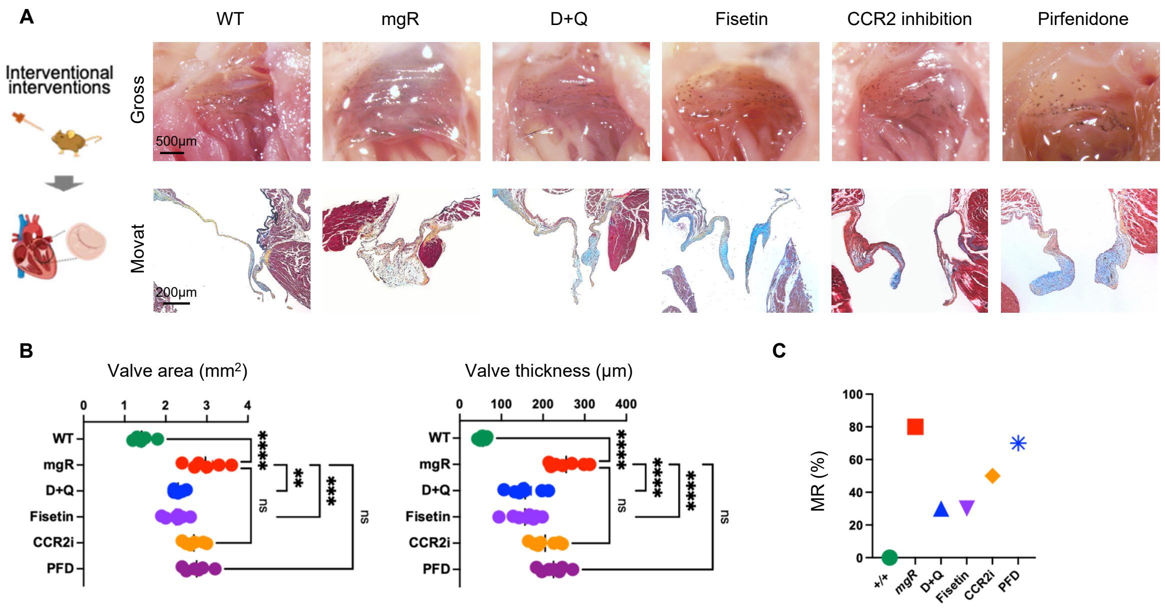


**Figure S6: Senolytic therapy restores ECM homeostasis and mitral valve function in MMVD**

**(A)** Gross morphology (upper) and histological features (lower) of the mitral valve across six experimental groups.

**(B)** Quantification of mitral valve area and leaflet thickness across the six experimental groups.

**(C)** Incidence of MR across the indicated experimental groups.

Data are represented as individual values with mean ± SEM; ns, not significant; **P < 0.01, ***P < 0.001, and ****P < 0.0001 by 1-way ANOVA.

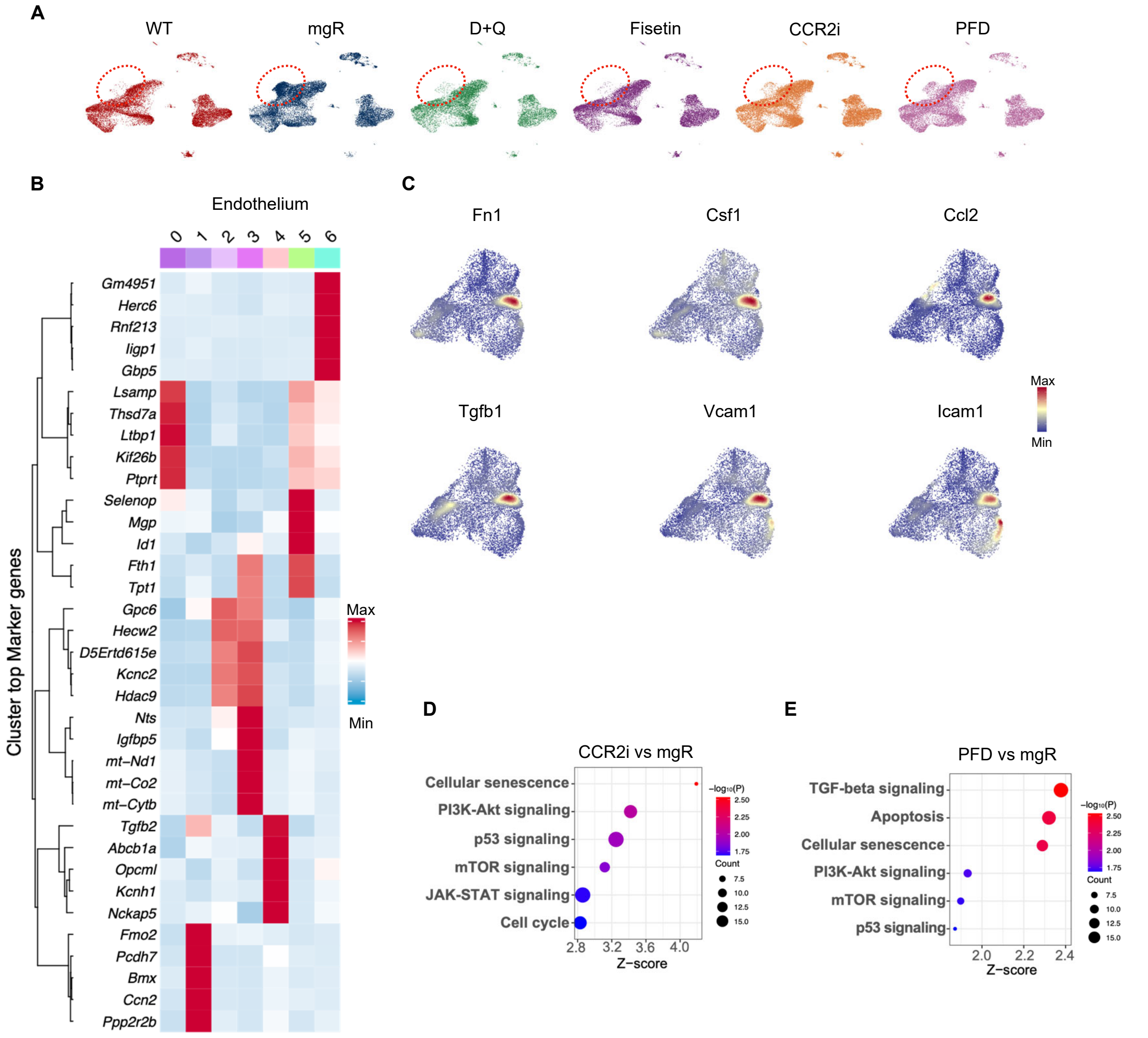


**Figure S7: Additional analyses of snRNA-seq data**

**(A)** UMAP of all mitral valve cells across the indicated experimental groups.

**(B)** Heatmap showing the top marker genes for each endothelial cluster.

**(C)** Expression of representative fibroinflammatory genes in endothelial cells.

**(D-E)** Bubble plots showing representative KEGG pathways enriched among DEGs in CCR2i vs mgR and PFD vs mgR comparisons.

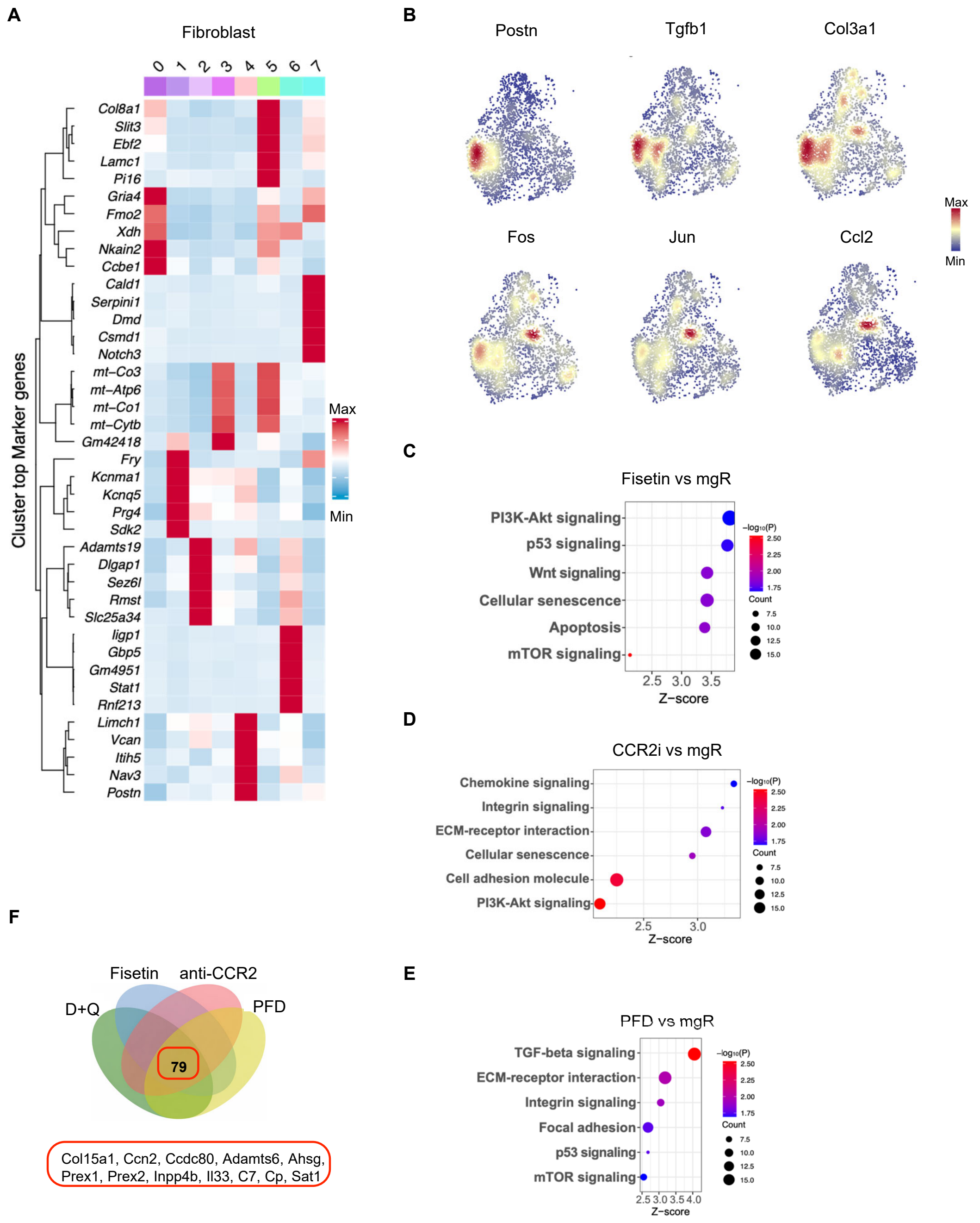


**Figure S8: Additional analyses of snRNA-seq data**

**(A)** Heatmap showing the top marker genes for each fibroblast cluster.

**(B)** Expression of representative fibroinflammatory genes in fibroblasts.

**(C-E)** Bubble plots showing representative enriched KEGG pathways derived from DEGs in fisetin versus mgR, CCR2i versus mgR, and PFD versus mgR comparisons.

**(F)** Venn diagram showing the overlap of DEGs from D+Q vs mgR, fisetin vs mgR, CCR2i vs mgR, and PFD vs mgR comparisons. A total of 79 genes are shared across all treatment groups.

**A**

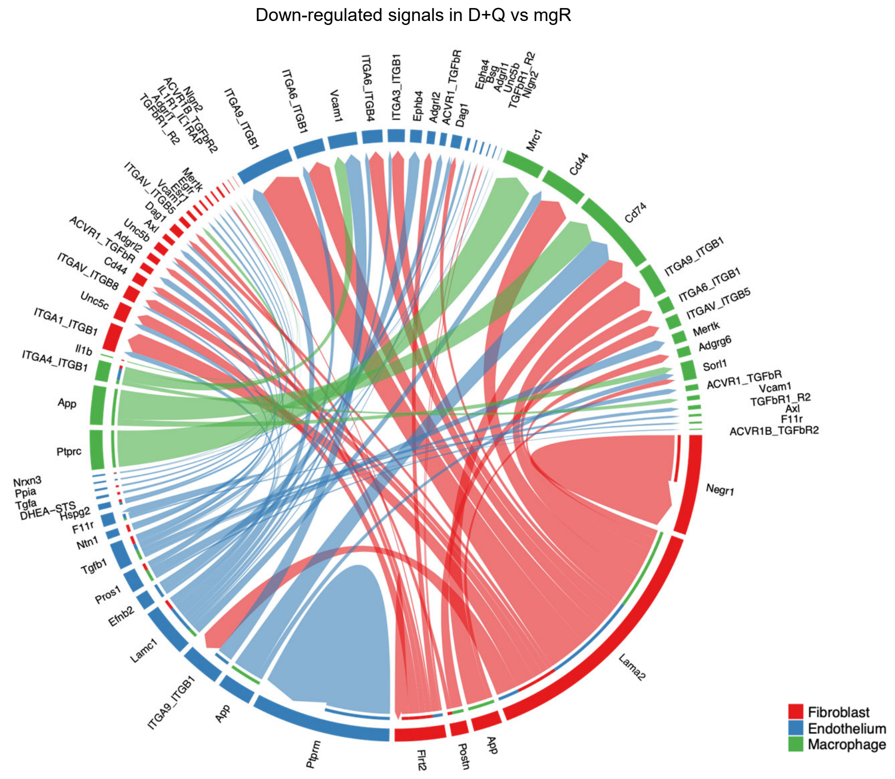


**B**

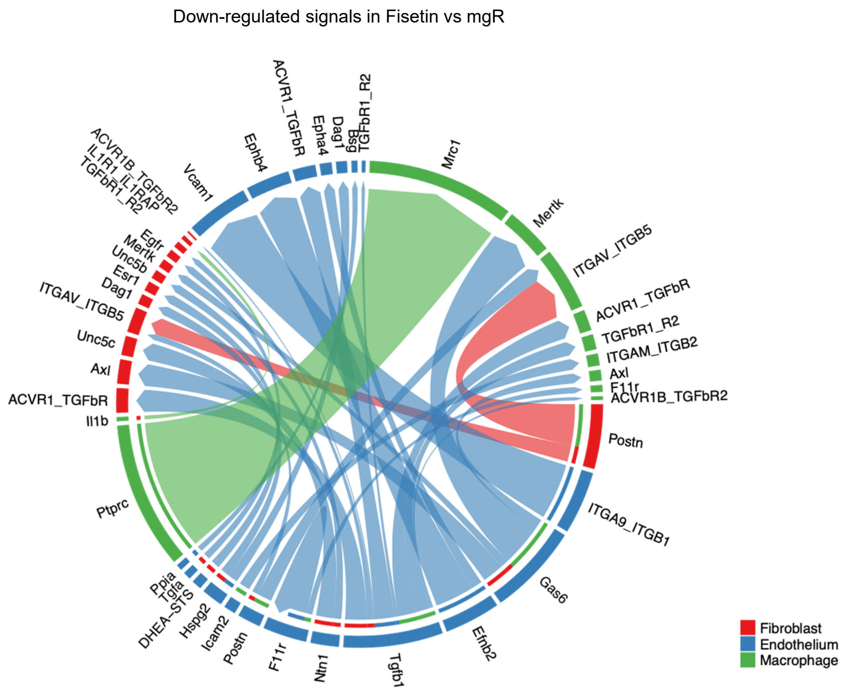


**Figure S9: Additional analyses of ligand-receptor interactions**

**(A)** Downregulated ligand-receptor interactions in D&Q versus mgR.

**(B)** Downregulated ligand-receptor interactions in fisetin versus mgR. Edge thickness reflects inferred interaction strength.

**A**

Down-regulated signals in CCR2i vs mgR

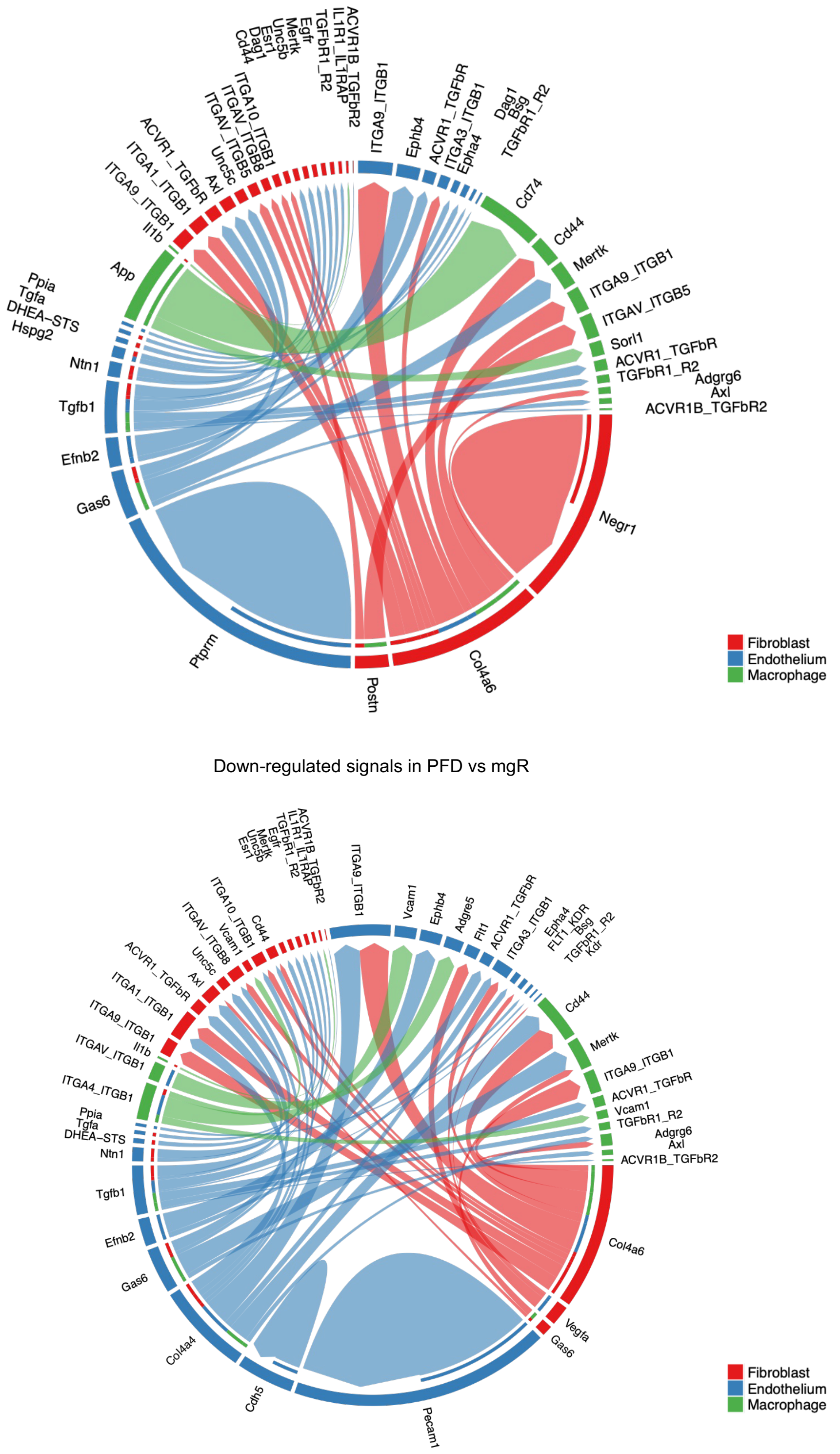


**Figure S10: Additional analyses of ligand-receptor interactions**

**(A)** Downregulated ligand-receptor interactions in CCR2i versus mgR.

**(B)** Downregulated ligand-receptor interactions in PFD versus mgR. Edge thickness reflects inferred interaction strength.

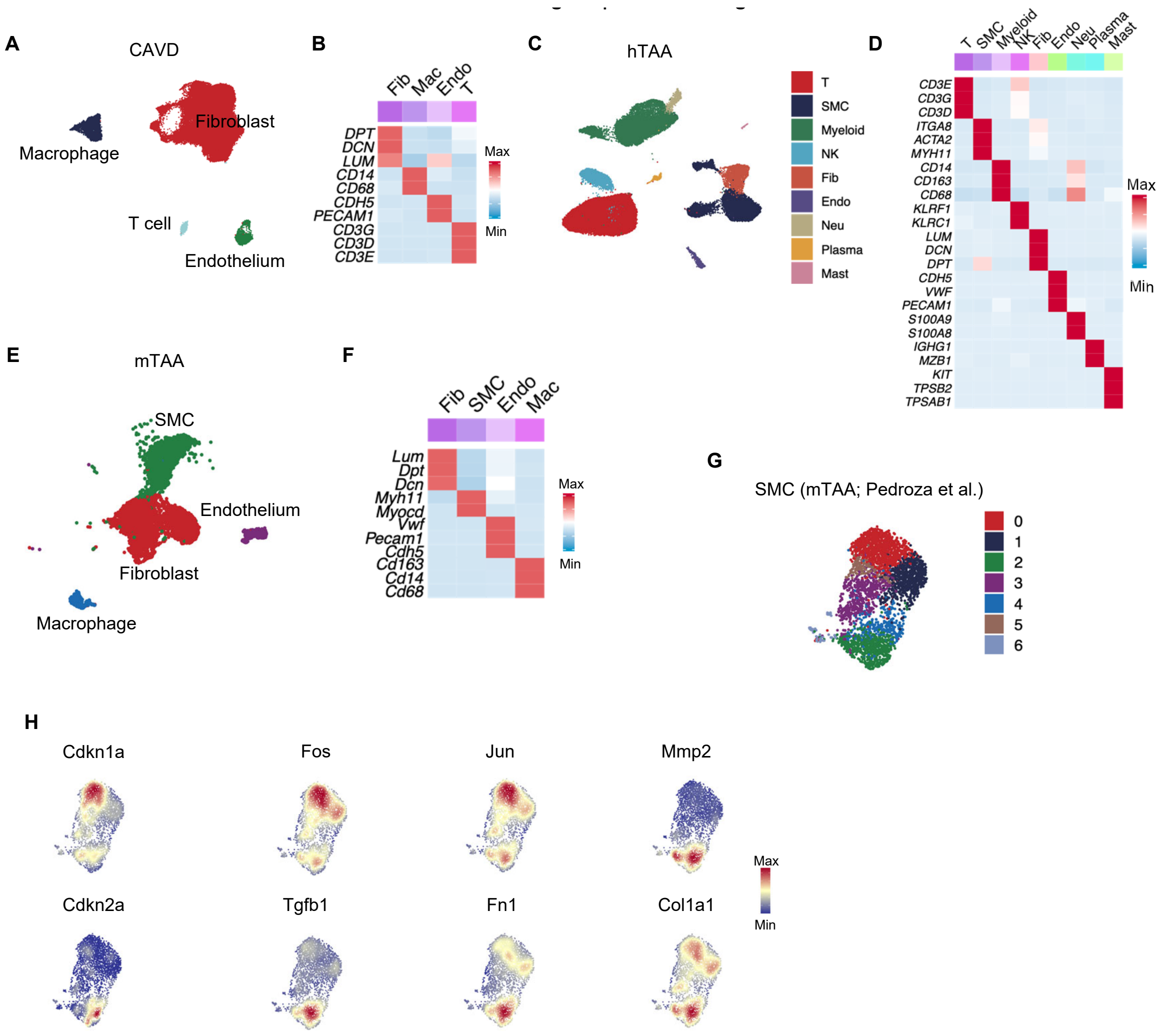


**Figure S11: Additional analyses of CAVD and TAA**

**(A)** UMAP showing cellular composition of CAVD.

**(B)** Heatmap showing the expression of canonical marker genes across major cell types in CAVD.

**(C)** UMAP showing cellular composition of hTAA.

**(D)** Heatmap showing the expression of canonical marker genes across major cell types in hTAA.

**(E)** UMAP showing the cellular composition of mouse thoracic aortic aneurysm (mTAA).

**(F)** Heatmap showing the expression of canonical marker genes across major cell types in mTAA.

**(G)** Unsupervised clustering of SMCs from mTAA.

**(H)** Expression of representative marker genes across SMCs from mTAA.
